# Semantic fragment representations for coordinate-free analysis of genomics data

**DOI:** 10.64898/2026.07.09.737627

**Authors:** Hamed Heydari, Jing Zhao, Madeleine Arseneault, Leila Younesian, Simon Tanguay, Yasser Riazalhosseini, Hani Goodarzi, Hamed Najafabadi

## Abstract

Many genomic assays begin with individual DNA fragments, but standard analysis quickly collapses those molecules into counts over genomic intervals. Rich information carried by each fragment, including its sequence, fragment body, cleavage boundaries, and local flanking context, is lost in this process. This loss is especially apparent in mixed-source and heterogeneous samples, where individual fragments originate from disparate cell types and can retain information about their cell of origin. To address this, we present LEAF-1, a fragment-level foundation model pre-trained on approximately 58 billion fragments spanning bulk ATAC-seq, single-cell ATAC-seq, and cell-free DNA profiles, representing each DNA molecule as a point in a learned semantic space defined by sequence context, assay modality, and explicit cleavage-boundary tokens. In sparse scATAC-seq datasets, mean-pooled LEAF-1 embeddings readily classify human cell types from as few as ∼1,000 fragments per cell, with high-scoring fragments linked to cell-type-associated transcription-factor programs. Similarly, in cell-free DNA profiling, LEAF-1 outperformed state-of-the-art coordinate-binning strategies and general-purpose DNA language model baselines across cancer detection tasks. Applying attention-based multiple-instance learning to LEAF-1 embeddings further improved cancer detection, reaching an area under the receiver operating characteristic (ROC) curve (AUC) of 0.95. This pan-cancer model generalizes beyond cancer types it is trained on, as we show by profiling plasma samples from clear cell renal cell carcinoma patients and healthy volunteers and applying the frozen classifier without retraining, achieving an AUC of 0.83. These results show that semantic learning over individual DNA fragments preserves biochemical, cell-associated, and disease-associated signals that are otherwise lost during coordinate-based aggregation.

## 1. Introduction

Analysis of DNA fragments forms the basis of most high-throughput genomics assays. For example, in chromatin-accessibility assays such as ATAC-seq, each fragment reflects a transposition event shaped by accessible regulatory DNA ^1^ and local sequence context ^2^. In cell-free DNA (cfDNA) profiling, each fragment reflects an in vivo cleavage event shaped by chromatin state in cell of origin, nuclease activity, and disease biology ^3;4^. In all cases, each fragment harbors rich information, including sequence, cleavage boundaries, and flanking context, that reflects the underlying experimental and biological conditions. Virtually all standard analysis frameworks, however, collapse this information early by aggregating fragments into genomic bins ^4;5^ or intervals (such as peaks ^6^) after alignment, and downstream models operate on the resulting count profile for almost all analytical goals. This representation has been effective, but it imposes a coordinate-based view that obscures biological relationships: related fragments map to distant loci and are treated as unrelated, while fragments from different cells of origin map to the same locus and are averaged together. This loss is especially consequential in sparse and mixed-source settings, where less-abundant phenomena become invisible through mapping to coordinates alone.

This information loss propagates in downstream analyses, including existing machine-learning (ML) approaches for various classification and prediction tasks. In scATAC-seq, for example, cell type identification models typically operate on peak-by-cell or region-by-cell matrices ^7–9^ or a transformation of such matrices ^10–13^. In cfDNA, fragmentomic methods such as DELFI ^5;14^, GALYFRE ^15^, EMIT ^16^, and related approaches augment interval-level counts with fragment length, end-position or motif features, but still summarize each sample into engineered aggregate features, discarding the joint structure of properties within each fragment and the ability to model fragments individually. We reasoned that DNA fragments should instead be modeled directly, where a sample can be viewed as a bag of fragments. By embedding these fragments into a learned shared semantic space, downstream analyses can operate on distributions of molecules in this semantic space rather than on their distribution in the genomic coordinate space. Unlike the coordinate space, in this semantic representation, proximity reflects shared biology, not simply chromosomal position, increasing information density (**Figure 1a**).

**Fig. 1.**
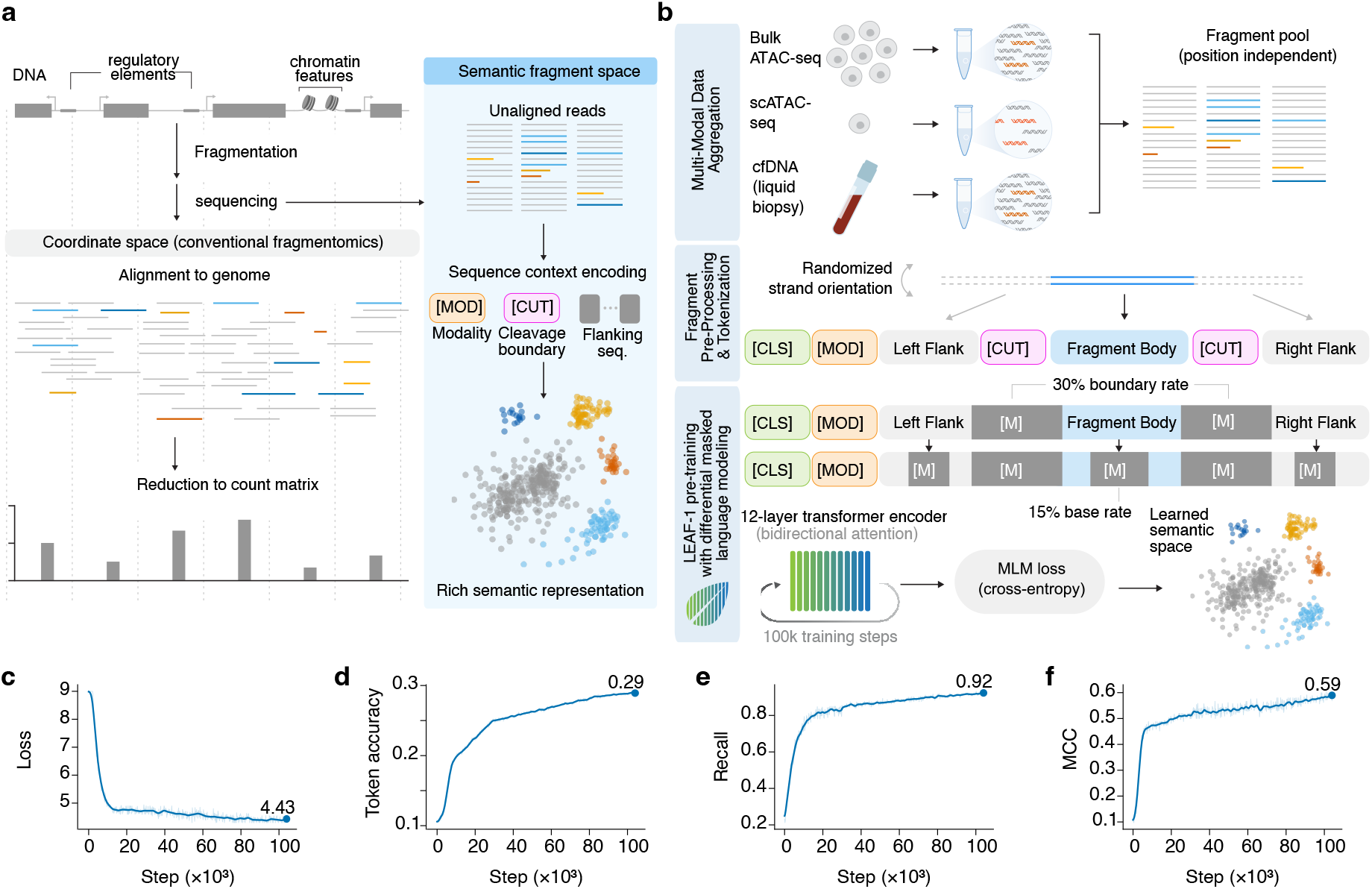
LEAF-1 learns semantic representations of DNA fragments. **(a)** Conventional fragmentomics analysis maps DNA fragments to genomic coordinates and aggregates them into fixed bins or predefined regions, producing a coordinate-count matrix. This representation treats fragments as related only when they overlap the same genomic interval. LEAF-1 instead embeds each fragment in semantic space using its sequence context, assay modality, fragment body, flanking sequence, and cleavage boundaries, so that fragments with shared biochemical or regulatory structure can be nearby even when they originate from distant genomic loci. **(b)** LEAF-1 pretraining workflow. Fragments from cell-free DNA, bulk ATAC-seq, and single-cell ATAC-seq are pooled into a position-independent corpus, tokenized as [CLS] [MOD] left flank [CUT] fragment body [CUT] right flank, and trained with masked language modeling. [CUT] boundary tokens are masked at higher frequency than sequence tokens and both boundaries are masked jointly, forcing the model to learn cleavage-associated sequence context. A 12-layer transformer encoder produces fragment embeddings used for downstream cell- and sample-level modeling. **(c)** Masked-language-model loss during pretraining. **(d)** Masked-token prediction accuracy. **(e)** [CUT] boundary recall. **(f)** [CUT] boundary Matthews correlation coefficient. Final values are shown at the end of training.

Here, we introduce LEAF-1 (learned embeddings for analysis of fragmentomics data), a genomic foundation model for semantic learning over DNA fragments. LEAF-1 explicitly encodes each fragment using its sequence context, assay modality, fragment body, local flanks, and explicit cleavage boundaries. We show that LEAF-1 embeddings can directly be used for downstream analytical tasks, bypassing coordinate-based binning altogether, including cell type identification from scATAC-seq data and cancer identification from cfDNA data, which represent two application cases with extreme sparsity and heterogeneity, respectively. LEAF-1 embeddings are biologically interpretable and, at the same time, substantially improve downstream ML performance over the established baselines, including in zero-shot identification of cancer types not included in its training data.

## 2. Results

### 2.1. LEAF-1 learns a semantic space for DNA fragments

LEAF-1 represents each DNA fragment as a learned point in semantic embedding space, defined by its sequence, cleavage boundaries, local flanking context, as well as assay chemistry rather than by genomic coordinates. This design encodes biological and experimental variables, including the regulatory context of the fragment (e.g., the cis-regulatory grammar dictating chromatin accessibility or nucleosome organization) and fragmentation biology (fragment end motifs) directly into the fragment embedding ^3;4;17^. Because different cells rely on different regulatory logic, a semantic space that captures this biology should place fragments from different cells in distinct regions, an organization we test directly in the sections that follow. Likewise, fragments carrying different fragmentomic features, such as the distinct end motifs of cfDNA released from healthy versus malignant cells, should fall in distinct regions of this space.

This representation required four design choices. First, the encoder operates on individual fragments rather than cell- or sample-level aggregates, preserving signals that would otherwise be diluted by averaging. Second, both cleavage boundaries are marked with explicit [CUT] tokens, allowing the model to learn sequence context around the fragment ends. Third, the fragment body and flanking sequence are retained, so that each embedding captures both the protected or accessible DNA segment and the local context that shaped its cleavage. Fourth, a modality token distinguishes Tn5-based from endogenous-nuclease fragmentation, the two major categories of fragmentation incorporated in our model, allowing one encoder to learn shared and chemistry-specific structure across assays.

LEAF-1 was pretrained on 58 billion fragments from 2,620 samples spanning cfDNA from healthy individuals and cancer patients ^18^, bulk ATAC-seq of tumors from different tissues ^19^, and single-cell ATAC-seq spanning different human cell types ^20^ (**Figure 1b & Supplementary Table 1**; see **Methods**). These modalities expose the model to two major fragmentation regimes: Tn5-mediated fragmentation in chromatin-accessibility assays and endogenous-nuclease fragmentation in plasma cfDNA. Each input followed the structure [CLS] [MOD] left_flank [CUT] fragment_body [CUT] right_flank within a 2,048-token budget, with dynamic flanking context and randomized strand orientation.

For pretraining, we used masked language modeling, with [CUT] tokens masked at twice the rate of ordinary sequence tokens. Both fragment boundaries were masked jointly, preventing the model from trivially inferring one boundary from the other and forcing prediction from the surrounding body and flank sequence. During training, masked-language-model loss decreased steadily (**Figure 1c**), with token accuracy reaching 0.29 (**Figure 1d**). Focusing on the [CUT]-token, prediction recall reached 0.92 (**Figure 1e**) with Matthews correlation coefficient (MCC) of 0.58 (**Figure 1f**). These results indicate that LEAF-1 learns cleavage-associated sequence structure rather than simply memorizing fragment position or length.

### 2.2. LEAF-1 embeddings preserve regulatory identity in single-cell ATAC-seq

To check whether the semantic fragment space learned by LEAF-1 preserved biologically useful signals in a downstream assay, we first examined the model’s ability to infer cell-type-associated regulatory programs from single-cell ATAC-seq data. To do this, we applied the frozen LEAF-1 encoder to single-cell ATAC-seq from 28 human tissues ^20^, where each cell is represented by a sparse set of Tn5-generated fragments. We down-sampled the data to test whether 1,000 fragments per cell are sufficient to recover cell identity without peak calling or coordinate-bin features. We also used these data to examine whether fragment-level classifier scores can be interpreted against known regulatory elements and transcription-factor programs. Each cell was represented by the mean of its frozen LEAF-1 fragment embeddings (**Figure 2a**). Using one-vs-all logistic regression across 108 cell types and 28 tissues, LEAF-1 embeddings reached mean AUC 0.93, median AUC ∼0.94, and ∼30-fold average-precision lift over prevalence (**Figure 2b,c** and **Supplementary Figures 1-3**; see **Methods**). Performance was highest for developmentally distinct lineage states (such as hepatocytes, pancreatic beta cells, and skeletal myocytes, AUC near 1.0) and lower for broadly defined or closely related cell types (such as endothelial subtypes and smooth muscle, AUC 0.70–0.82), consistent with embeddings capturing regulatory identity rather than technical artifacts such as sequencing depth or batch effects. A UMAP of mean-pooled cell embeddings showed corresponding organization by cell-type category (**Figure 2d**).

**Fig. 2.**
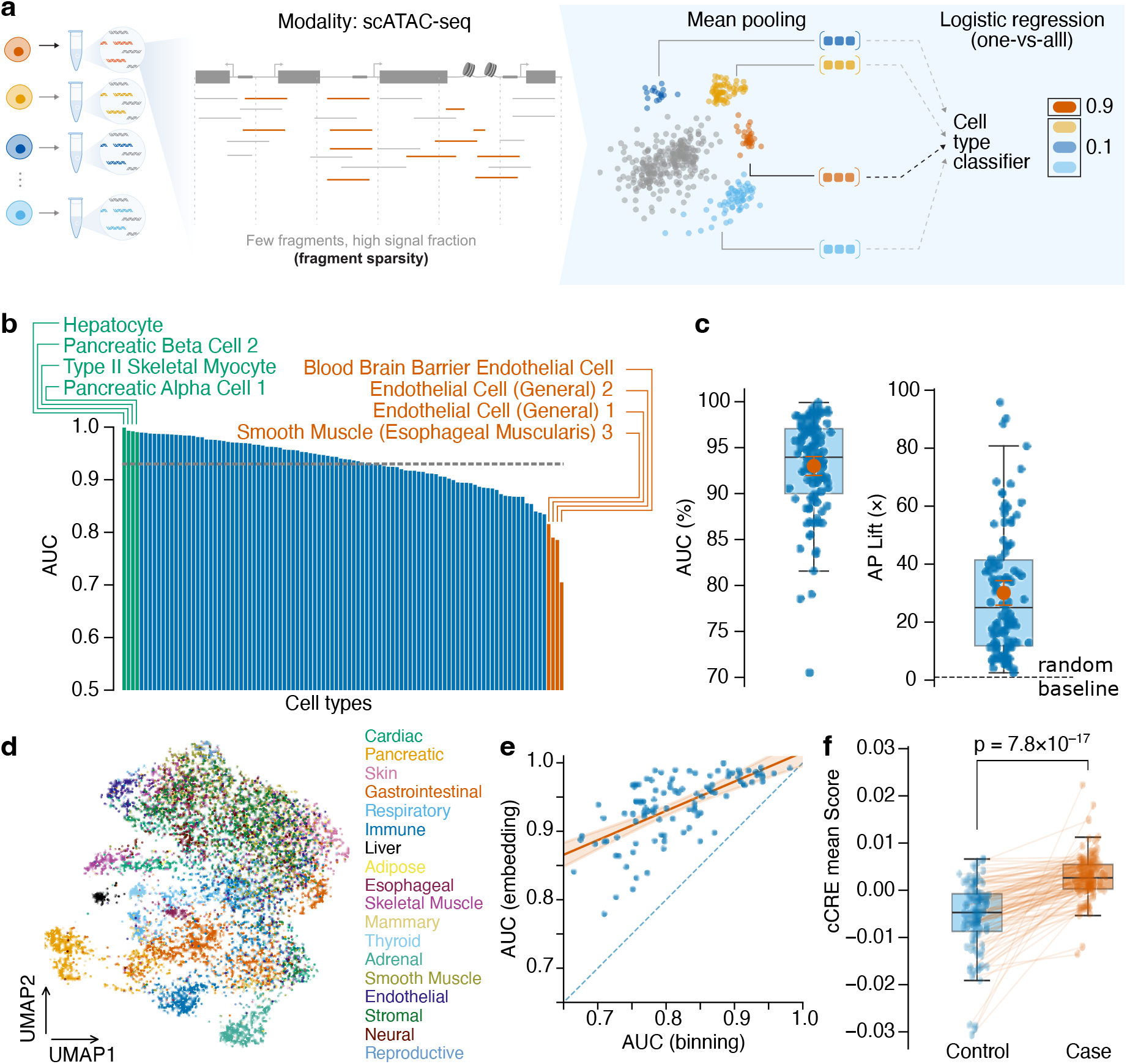
LEAF-1 embeddings classify single-cell ATAC-seq cell types and localize informative fragments to regulatory elements. **(a)** Schematic of the cell-level classification approach. Each cell’s fragments are individually embedded by the frozen LEAF-1 encoder and averaged into a single mean-pooled cell embedding, which is passed to a one-vs-all logistic-regression classifier to predict cell type across 108 cell types from 28 human tissues. **(b)** One-vs-all classification AUC for 108 cell types across 28 tissues using mean-pooled LEAF-1 embeddings from the frozen encoder. Cell types are ranked by AUC. Green labels indicate top-performing cell types, including hepatocyte, pancreatic beta cell 2, type II skeletal myocyte, and pancreatic alpha cell 1; orange labels indicate lower-performing or broadly defined cell types, including endothelial cell subtypes, smooth muscle, naive T cell, and blood-brain barrier endothelial cell. Dashed line, mean AUC; shaded band, interquartile range. **(c)** Distribution of one-vs-all AUC values across cell types, shown as percent AUC, and average-precision lift over the random baseline. **(d)** UMAP projection of mean-pooled cell embeddings, colored by annotated cell type, showing organization of cells in LEAF-1 embedding space. **(e)** Per-cell-type AUC for LEAF-1 embeddings versus coordinate-binning features under matched evaluation. Each point is one cell type; dashed diagonal indicates equal performance; orange line indicates fitted trend with 95% confidence band. **(f)** Fragment-level classifier scores at cell type’s associated candidate cis-regulatory elements (cCREs). For these cCREs, scores are higher in the matching cell type (Case) than in control cell types (Control) (paired two-sided t-test across n = 102 cell types, t = 10, P = 7.8 × 10^−17^), indicating that high-scoring LEAF-1 fragments localize to cell type-relevant regulatory elements.

We specifically opted for representing each cell with the mean of its fragment embeddings rather than more complex aggregation strategies such as set-based representation learning since this approach mirrors interval-based representation, allowing us to directly compare the semantic and coordinate spaces. Specifically, interval-based count aggregation (followed by depth normalization per cell) is equivalent to representing each fragment with a one-hot-encoded vector that specifies the interval it belongs to, followed by calculating the mean of these one-hot-encoded embeddings. Thus, we reasoned that any difference in downstream performance using semantic mean embeddings vs interval-based counts represents the information density of the semantic vs coordinate space. We compared LEAF-1 embeddings with coordinate-bin representations under matched cross-validation, and observed that per-cell-type AUCs for LEAF-1 matched or exceeded coordinate binning across all cell types (AUC gain +0.01 to +0.24), with the largest gains in cell types where binning performed weakest (AUC < 0.75) (**Figure 2e** and **Supplementary Figure 4**). Because the encoder was frozen and the downstream classifier was held fixed, this comparison isolates the representation supplied to the classifier, indicating that fragment embeddings recover signals that coordinate counts capture less effectively, with the largest gains where coordinate binning performs weakest.

We then asked where the classifier’s cell type-distinguishing ability comes from. If the classifier is biologically grounded, its signal should come from fragments in regulatory elements active in the matching cell type. To test this, we scored each fragment individually rather than using the per-cell mean. For each of 102 cell types, we compared, for fragments originating from that cell type’s associated candidate cis-regulatory elements (cCREs) ^20^, the scores assigned by each cell type-matched model to its corresponding cell type versus the scores it assigned to other cell types (**Methods**). Scores at these cCREs were higher in the matching cell type (paired two-sided t-test across n = 102 cell types, t = 10, P = 7.8 × 10^−17^; **Figure 2f**). The classifier’s ability to distinguish cell types therefore comes largely from fragments in cell type-specific regulatory elements.

We also observed that fragment-level scores reflect transcription-factor programs (**Figure 3**). Specifically, within individual cell types, we used FIRE ^21^ to compute a motif enrichment statistic based on mutual information between motif occurrence and fragment score, producing a specific enrichment value for each TF in each of the 102 one-vs-all comparisons. Positive values indicate that a motif predominantly characterizes highly scored fragments, whereas negative values indicate an association with lowly scored fragments. By this measure, fragment level scores correlated significantly with the presence of established TF motifs (**Figure 3a**). For example, in club cells, FOXA2 motif enrichment was tightly correlated with the mean LEAF-1 score across score-ranked fragment bins (Pearson r = 0.92; **Figure 3b**). This is consistent with the established role of FOXA2 as a pioneer transcription factor in airway epithelium, where it directly regulates club cell–specific genes and chromatin accessibility ^22^. This robust association suggests that LEAF-1 scores capture authentic regulatory context and TF binding sequence rather than confounding baseline properties such as GC content or fragment length.

**Fig. 3.**
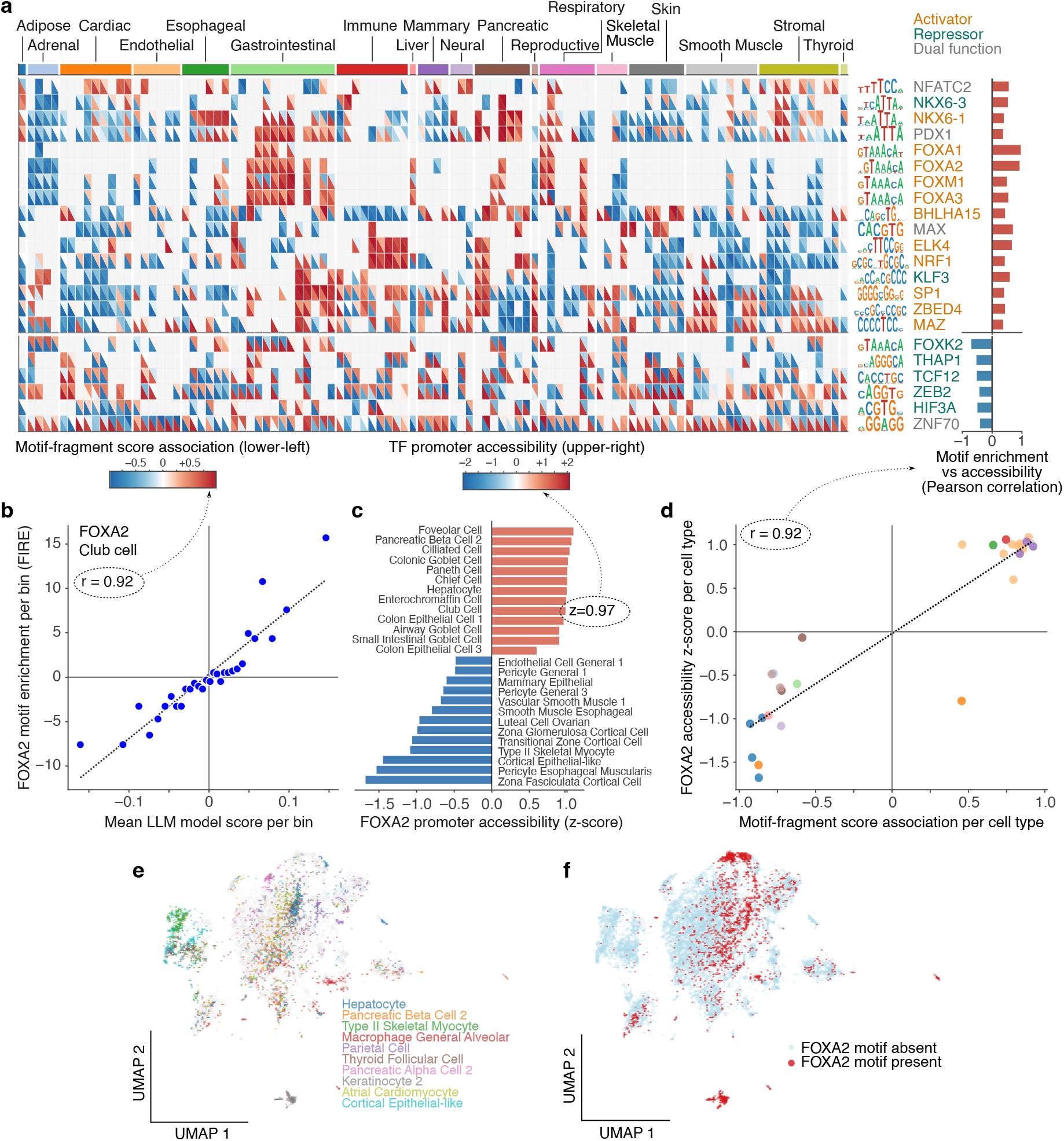
LEAF-1 fragment scores recover transcription-factor programs in single-cell ATAC-seq. **(a)** Heatmap of significant transcription-factor motif associations across cell-type comparisons. Rows are TF motifs, with sequence logos shown at right. Columns are one-vs-all cell-type comparisons, annotated by tissue category in the top color bar. Each heatmap cell is split diagonally: the lower-left triangle shows the association between LEAF-1 fragment score and motif enrichment, and the upper-right triangle shows the promoter chromatin accessibility of the corresponding TF in the same cell type. Red indicates positive association or increased accessibility; blue indicates negative association or decreased accessibility. Motifs are grouped into activating-enriched factors with positive score-accessibility concordance (top block) and repressive-enriched factors with negative or context-repressive concordance (bottom block); the color of each TF at right denotes its annotated role (activator, repressor, or dual) based on known TF function. The association between TF function and motif-accessibility direction is assessed by Fisher’s two-sided exact test (n = 18, p = 0.0025). Right bars summarize the overall motif-enrichment and accessibility correlation per TF. **(b)** Per-bin relationship underlying the lower-left triangle value, shown for FOXA2 in Club Cell: mean motif enrichment versus mean LEAF-1 model score across 30 bins (Pearson r = 0.92), with points colored by mean model score. **(c)** Derivation of the upper-right triangle value for FOXA2: promoter TN5 insertion counts are converted to pseudobulk log-CPM and then z-scored across cell types, giving the accessibility value shown in the heatmap. **(d)** Concordance between the two triangle values across 27 cell types for FOXA2: motif enrichment score (lower-left triangle) versus promoter accessibility z-score (upper-right triangle), colored by tissue category (bar-chart r = 0.92). **(e)** Each fragment is scored by the model across cell types, and these per-cell-type scores are projected into the semantic regulatory space. UMAP computed on per-cell-type fragment scores (fragment 102 cell-type scores). Fragments are colored by cell type, showing the ten cell types with highest AUC, remaining fragments are shown in light gray. Fragments organize into regions corresponding to distinct cell-type programs. **(f)** The same UMAP colored by FOXA2 motif presence: fragments where the motif is present (red) or absent (light blue). FOXA2 containing fragments concentrate in specific neighborhoods of the embedding rather than distributing uniformly.

Moreover, across cell type comparisons, the direction of these associations was consistent with the established regulatory roles of the respective factors. For every TF evaluated, we computed the Pearson correlation between its motif enrichment and its promoter accessibility across the 102 comparisons (**Figure 3c**,**d**; see **Methods**). TFs are separated into two broad groups according to the sign of this correlation (**Figure 3a**; right panel). Several factors with predominantly activating regulatory functions, including FOXA family members such as FOXA2 (**Figure 3d**), as well as NKX6-1, BHLHA15, ELK4, NRF1, SP1, ZBED4 and MAZ, displayed positive correlations. Specifically, their motifs were enriched within highly scored fragments in the same cell types where the TF’s promoters were most accessible. Conversely, repressive or context repressive factors, including FOXK2, THAP1, TCF12, ZEB2, HIF3A, and ZNF70, displayed the inverse pattern, with their motifs enriched in cell types where the TF promoter accessibility was lower (two-sided Fisher’s exact test for association between TF function and motif-accessibility direction: n = 18 activator- or repressor-annotated TFs, P=0.0025).

We note that, given the linear nature of our cell type-scoring, each one-vs-all classifier represents a constant vector in the LEAF-1 embedding space. Thus, cell type scores effectively represent projections of the embeddings onto these vectors. **Figure 3e** visualizes this organization at the fragment level, showing the UMAP of the fragment embeddings after projection onto cell type vectors. As expected, fragments separate into regions corresponding to distinct cell type-specific programs (**Figure 3e**). For example, FOXA2-containing fragments concentrate in specific neighborhoods of this embedding rather than distributing uniformly (**Figure 3f**), consistent with the motif-level associations above.

### 2.3. Boundary-aware fragment representations improve disease detection from mixed-source cfDNA data

We next evaluated LEAF-1 performance to detect cancer signatures in cfDNA data, a mixed-source setting in which fragment-level modeling should be especially useful. Plasma cfDNA contains molecules released from many tissues across the body, predominantly hematopoietic cells ^23^, as well as diseased cell-associated fragments that often represent only a small minority of the total sample. In contrast to Tn5-generated scATAC fragments, cfDNA fragments are produced by endogenous nucleases, including DNASE1L3, DNASE1, and DFFB, whose cleavage preferences are shaped by sequence and chromatin context ^3;4;17;24^. cfDNA, therefore, provides a pivotal test of the ability of LEAF-1’s boundary-aware fragment representations to better recover disease-associated signals relative to interval-based approaches (**Figure 4a**).

**Fig. 4.**
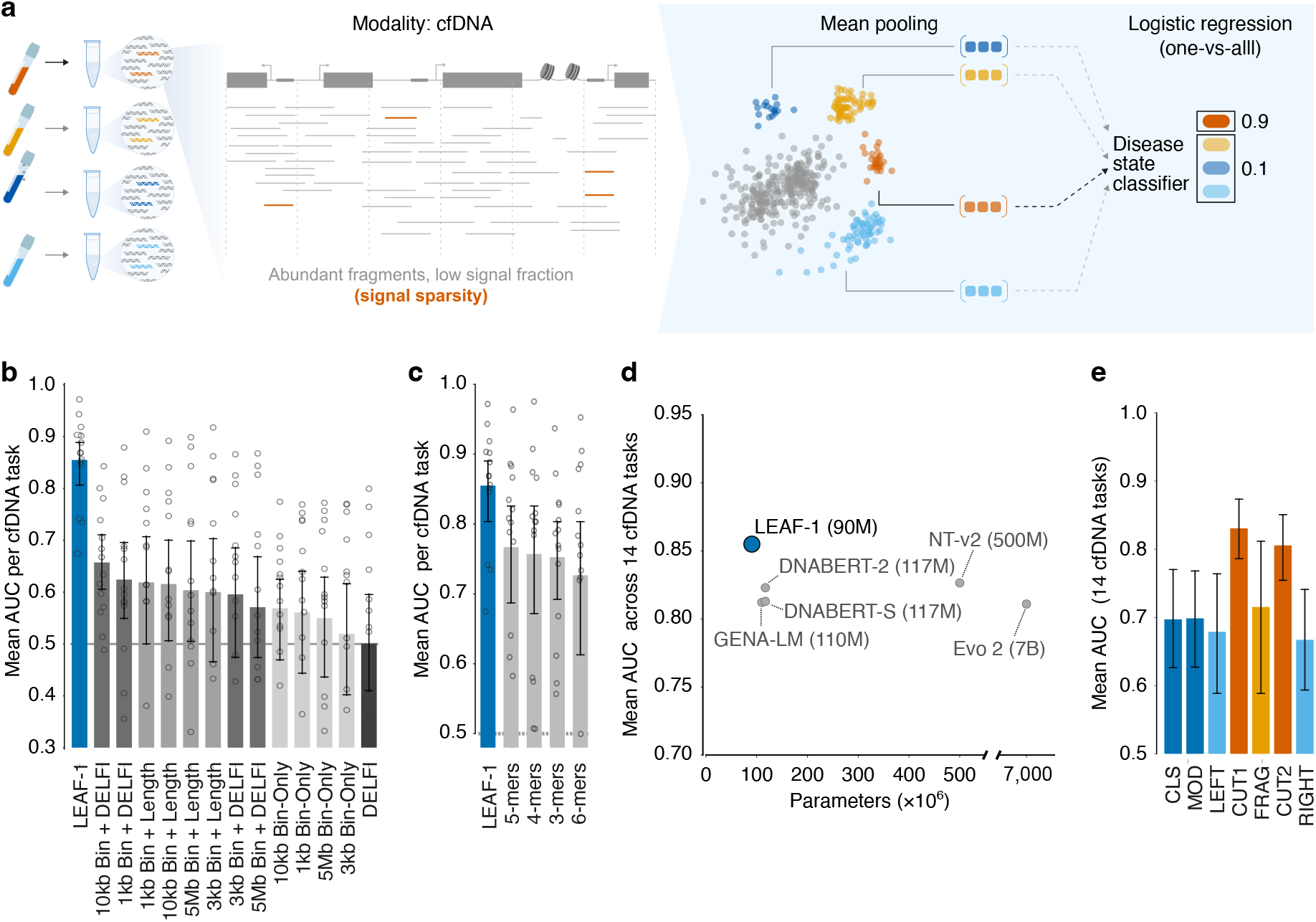
LEAF-1 outperforms coordinate, sequence-composition, and DNA language-model baselines in cf DNA, with signal concentrated at cleavage boundaries. **(a)** Schematic of the cf DNA-based disease state classification approach. cf DNA fragments extracted from plasma samples are embedded by the frozen LEAF-1 encoder and mean-pooled into a single sample-level embedding, which is passed to a one-vs-all logistic-regression classifier to predict disease state. All panels evaluate frozen LEAF-1 embeddings under matched leave-one-out nested cross-validation across 14 cf DNA cancer-detection tasks, with each sample fixed to 10,000 fragments and linear classifiers used for downstream prediction. **(b)** Mean AUC for LEAF-1 compared with coordinate-binning and DELFI-style fragmentomic baselines. Baselines include Bin-only, Bin+Length, and Bin+DELFI features across multiple bin resolutions, plus DELFI alone. Dots show task-level AUCs; bars show mean AUC; error bars indicate 95% confidence intervals. Dashed line, chance. **(c)** Mean AUC for LEAF-1 compared with canonical k-mer frequency baselines from k = 3 to 6. LEAF-1 improved mean AUC over all k-mer baselines, with 95% bootstrap confidence intervals for paired AUC differences excluding zero. **(d)** Mean AUC versus model size for LEAF-1 and external DNA language-model baselines, including GENA-LM, DNABERT-2, DNABERT-S, NT-v2, and Evo 2, evaluated under the same matched protocol. **(e)** Single-segment analysis of LEAF-1 input components. Classifiers trained on individual segment embeddings show that cf DNA signal is strongest at the two cleavage-boundary tokens (CUT1 and CUT2), exceeding the fragment body, flanking sequence, and special-token embeddings. Bars show mean AUC across 14 tasks; error bars indicate 95% confidence intervals.

Using cf DNA data from Cristiano et. al ^5^, comprising 538 samples from seven cancer types and healthy controls, we first benchmarked the frozen representation using mean-pooling followed by logistic regression for 14 cancer detection tasks; this includes seven cancer vs healthy and seven cancer vs other cancers (tissue-of-origin) binary classification tasks. We down-sampled the data to 10,000 fragments, and linear classifiers were trained under matched leave-one-out nested cross-validation so that all methods used the same fragments and differed only in representation (Refer to **Supplementary Figure 6** for a schematic diagram of the process). Across all cfDNA cancer-detection tasks, LEAF-1 exceeded coordinate-binning and DELFI-style baselines by +0.198 to +0.354 mean AUC (+30% to +71% relative) (**Figure 4b**; see **Methods**). These baselines included bin-only and bin+fragment length features at multiple resolutions, DELFI-style fragmentation features, and bin+DELFI combinations, allowing direct comparison against both interval-count and engineered fragmentomic summaries (see **Methods** for details).

LEAF-1 also outperformed direct sequence-composition baselines. Specifically, compared with k-mer frequency vectors (enumerating the number of times each k-mer is seen across all fragments; k = 3 to 6), LEAF-1 improved mean AUC by +0.088 to +0.128 (**Figure 4c**), with 95% bootstrap confidence intervals for the paired AUC differences always excluding zero (**Supplementary Figure 7a**). We also compared LEAF-1 against five general-purpose DNA language models, DNABERT-2 ^25^, DNABERT-S ^26^, GENA-LM ^27^, NT-v2 ^28^, and Evo 2^29^. LEAF-1 improved mean AUC by +0.029 to +0.044 under the same matched protocol (**Figure 4d**), by scale alone with 95% bootstrap CIs for the paired differences again excluding zero (**Supplementary Figure 7b**).

Importantly, this improvement over general-purpose DNA models was obtained despite the fact that LEAF-1 was the smallest model in this comparison, suggesting that fragment-specific pretraining and explicit cleavage boundary modeling provide information not captured by scale alone. Supporting this notion, token importance analysis localized the cfDNA signal used for cancer detection to fragment boundaries. Specifically, classifiers trained on the two [CUT] token embeddings reached AUC 0.831 and 0.806, outperforming the fragment sequence alone (0.715) and flanking sequence alone (∼0.67; **Figure 4e**). This pattern contrasts with scATAC, where signal was concentrated in the fragment body (**Supplementary Figure 5**), and is consistent with the biology of plasma DNA fragmentation, wherein endogenous nucleases leave sequence-dependent cleavage signatures at fragment ends ^4^.

### 2.4. Attention-based pooling improves pan-cancer detection

We next asked whether learned pooling could improve sample-level cf DNA prediction beyond mean-pooled fragment embeddings. Because cf DNA samples have sample-level labels but contain many unlabeled fragments, this setting naturally fits the application of multiple-instance learning (MIL): each plasma sample is treated as a bag of fragment instances, and the model learns how to combine their embeddings into one prediction. We implemented this as LEAF-1+MIL, pairing the frozen LEAF-1 encoder with a gated-attention MIL head ^30^; (**Figure 5a**; see **Methods**). Mean pooling assumes that all fragments contribute equally, which can dilute rare tumor-associated molecules into the background of normal cell turnover. In contrast, attention-based MIL learns fragment weights and can upweight informative regions of the fragment distribution while keeping the encoder frozen. To avoid collapse of the pooled representation when attention was broadly distributed across many fragments, we used effective-sample-size rescaling of the attention-weighted mean; implementation details are provided in **Methods**.

**Fig. 5.**
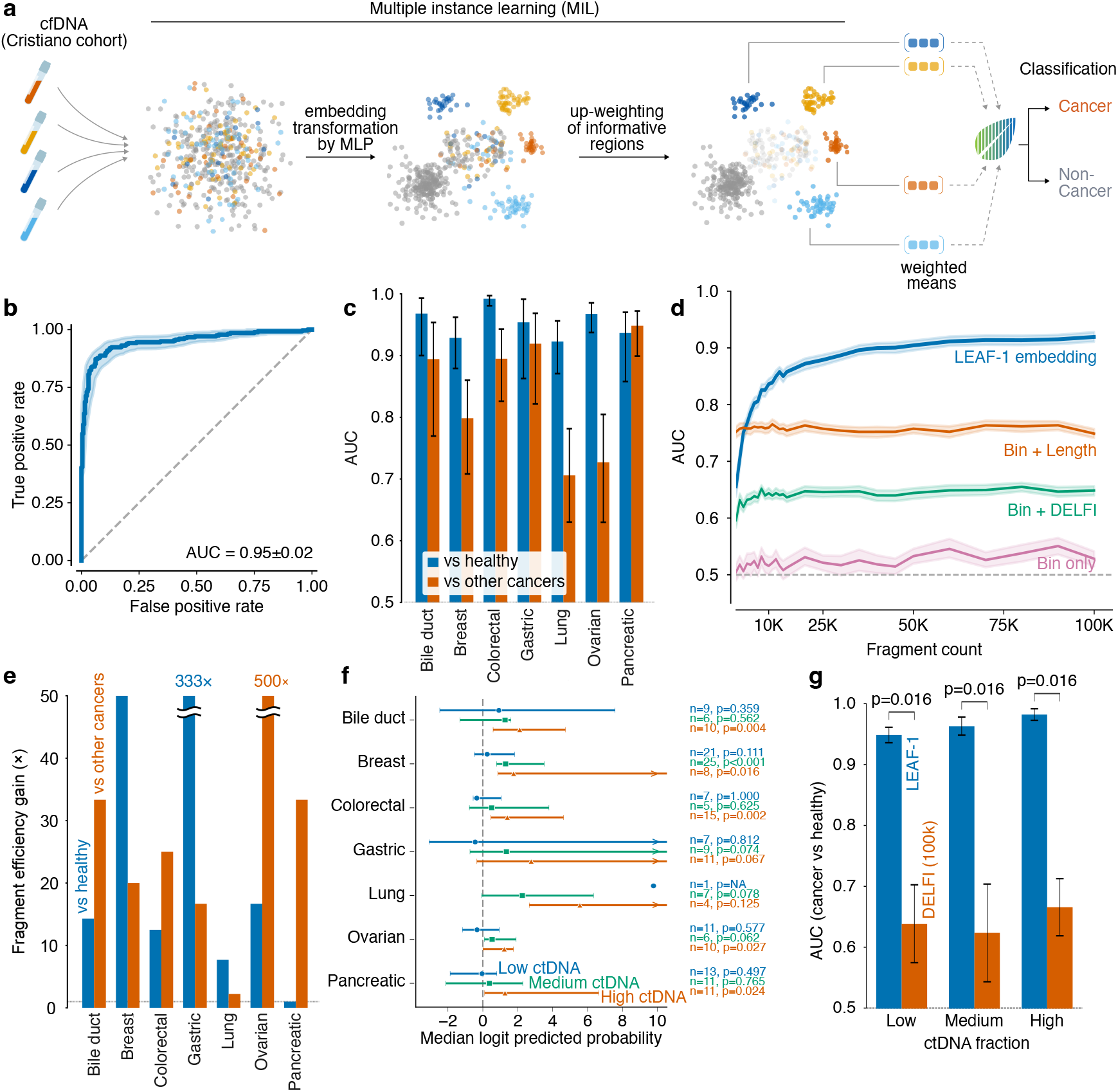
LEAF-1+MIL improves cf DNA cancer detection and fragment efficiency. **(a)** Schematic of the cf DNA classification approach based on multiple-instance-learning (MIL). Each cf DNA sample is treated as a bag of fragment instances whose frozen LEAF-1 embeddings are transformed by an MLP; a gated-attention head then up-weights informative regions of the fragment distribution and combines them into attention-weighted means, yielding a single sample-level representation that is classified as cancer or non-cancer. **(b)** Pan-cancer receiver-operating-characteristic curve for cancer-versus-healthy classification in the Cristiano cohort using LEAF-1+MIL with 100,000 fragments per sample. The model reached AUC 0.95 (95% CI, 0.93-0.97); shaded band indicates bootstrap confidence interval, and dashed diagonal indicates chance. **(c)** Per-cancer performance across seven tumor types. Blue bars show cancer-versus-healthy detection; orange bars show discrimination of each cancer type from the other cancer types. Error bars indicate 95% bootstrap confidence intervals; dashed line indicates chance. **(d)** Pan-cancer AUC as a function of the number of fragments sampled per sample. LEAF-1+MIL embeddings improve with fragment count and outperform coordinate baselines across the full range from 1,000 to 100,000 fragments. Shaded bands indicate confidence intervals; dashed line indicates chance. **(e)** Fragment-efficiency gain by cancer type for detection against healthy controls and discrimination against other cancers. Gain is defined as the fold reduction in fragments required for LEAF-1+MIL to match the coordinate-binning AUC at 100,000 fragments. Off-axis values are annotated. **(f)** Forest plot of LEAF-1 predicted probability (positive samples vs. healthy) across cancer types, split by ctDNA group. Predicted probabilities are logit transformed, points show the median logit per cancer and group, and bars show the 95% bootstrap confidence interval of the median. The dashed line marks logit(0.5) = 0, per group sample sizes and p-values (two-sided one sample Wilcoxon signed-rank test against 0) are annotated at right. Arrows indicate points or intervals extending beyond the axis limit. Higher ctDNA burden is associated with stronger LEAF-1 predictions. **(g)** AUC for LEAF-1 versus DELFI (100k features) within each ctDNA group. Bars show the mean AUC across the seven cancer types and error bars show the standard error of the mean (SEM). LEAF-1 shows a consistently higher AUC than DELFI across all three burden levels (two-sided paired Wilcoxon signed-rank test across cancer types, n=7, p = 0.016 for each comparison).

On the Cristiano et al cohort ^5^ used in the previous section, LEAF-1+MIL with 100,000 fragments per sample reached pan-cancer detection AUC of 0.949 (95% CI 0.93-0.97; 10,000-resample bootstrap; **Figure 5b**). Per cancer type, cancer-vs-healthy detection AUCs ranged from 0.992 for colorectal cancer to 0.923 for lung cancer, with bile duct, ovarian, gastric, pancreatic, and breast cancers reaching 0.968, 0.967, 0.954, 0.937, and 0.929, respectively (**Figure 5c** and **Supplementary Figure 8**). Multi-cancer discrimination, i.e., distinguishing one cancer type from others, was strongest for pancreatic, gastric, and colorectal cancer, and weakest for lung and ovarian cancer, suggesting weaker tissue-of-origin separation for the latter types under this protocol (**Figure 5c)**.

LEAF-1+MIL outperformed coordinate-based approaches across the three successively more informative baselines, Bin-only, Bin+Length, and Bin+DELFI, at all examined bin sizes (10K-300K), both for distinguishing cancer from healthy and for classifying tissues of origin (**Supplementary Figures 9-11**). The largest performance gain was observed over the least complex Bin-only features. Notably, this advantage persisted even when LEAF-1+MIL was compared to the best DELFI-informed baseline, and finer binning did not enable coordinate models to close the gap. We also tested fragment efficiency by measuring a series of subsamples ranging from 1,000 to 100,000 fragments and then performing the same pan-cancer analysis. LEAF-1+MIL reached pan-cancer AUC ∼0.90 by 20,000 fragments and ∼0.93 at 100,000 fragments, whereas the strongest coordinate baseline plateaued near 0.76 (**Figure 5d** and **Supplementary Figures 12-13**). The corresponding fragment-efficiency gains were largest for gastric cancer detection and ovarian cancer discrimination (**Figure 5e**), indicating that semantic fragment embeddings can recover cancer-associated signals from substantially fewer molecules than interval-count features.

To investigate whether LEAF-1 performance is associated with circulating tumor DNA (ctDNA) burden, we used ichorCNA-estimated tumor fraction ^31^ as a proxy for ctDNA burden and stratified cancer samples into low, medium, and high tumor fraction groups. LEAF-1-predicted probabilities increased with tumor fraction across cancer types, indicating that samples with higher inferred ctDNA burden tended to receive more confident positive predictions (**Figure 5f**). We then evaluated case-control discrimination within each tumor fraction group by comparing cancer samples in that group against healthy controls. LEAF-1 consistently outperformed DELFI across all three groups, achieving AUCs of approximately 0.94–0.98 compared with 0.62–0.67 for DELFI. This improvement was significant in each tumor fraction group (two-sided paired Wilcoxon signed-rank test on matched cancer-type AUCs for cancer-versus-healthy classification between the two methods, n = 7 cancer types, P = 0.016 for each group; **Figure 5g**). Notably, LEAF-1 maintained strong discrimination even in the low tumor fraction group, where ctDNA-derived signal is weakest, suggesting that LEAF-1 is robust for detecting cancers with low ctDNA burden.

### 2.5. LEAF-1 generalizes beyond cancer types in its training set

We next asked whether a pre-trained pan-cancer LEAF-1+MIL classifier could detect a cancer type not used during training. To address this question, we isolated and performed shallow whole-genome-sequencing of plasma cfDNA samples from 6 healthy donors and 6 patients with clear cell renal cell carcinoma (ccRCC) (**Supplementary Table 2**, see **Methods**), and then applied the model trained on seven cancer types in the Cristiano et al cohort (previous section) without retraining (**Figure 6a**). We obtained ∼853,000 to ∼1,587,000 fragment sequences per sample (median ∼957,000), corresponding to an ultra-low-pass whole-genome coverage of ∼0.1×. ichorCNA ^32^ analysis showed flat genome-wide copy-number profiles in both cancer patients and healthy donors, with log_2_ ratios tightly distributed around zero across all autosomes (**Figure 6b**). For the cancer samples, this indicates that the ctDNA fraction was insufficient for reliable detection of copy number alterations at this sequencing depth (a no-CNA profile is expected for healthy donors).

**Fig. 6.**
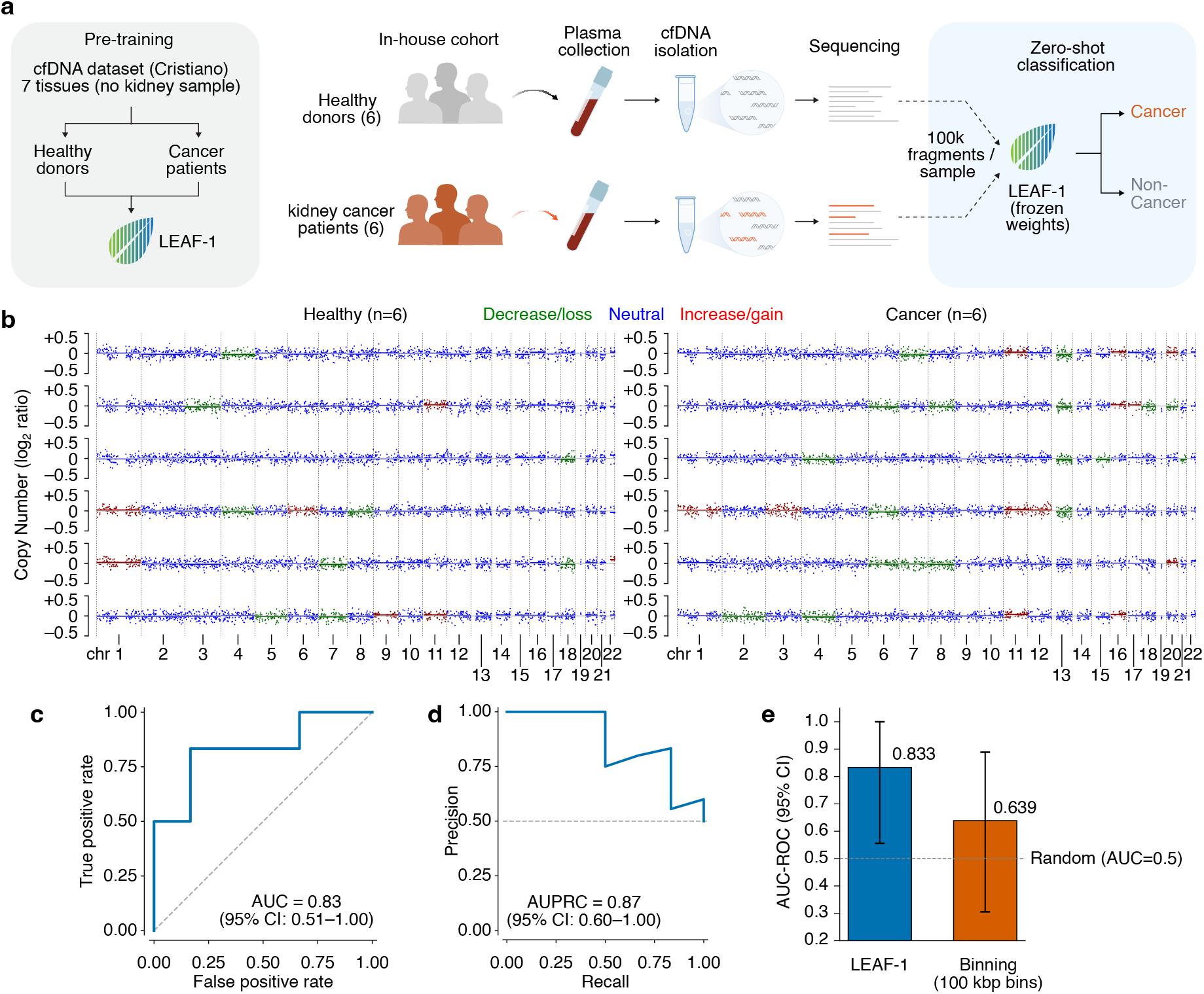
Zero-shot cf DNA detection of kidney cancer in an independently profiled in-house cohort. **(a)** Schematic of the analysis. A frozen pan-cancer LEAF-1+MIL classifier trained on the seven Cristiano cancer types was applied without retraining or threshold adjustment to 12 independently profiled plasma cfDNA samples: six healthy volunteers and six patients with kidney cancer, a cancer type absent from supervised model development. **(b)** Genome-wide copy-number profiles for the six healthy (left) and six kidney cancer (right) samples. Log_2_ copy number ratios across chromosomes 1–22 are shown; individual bins and copy number segments are colored by ichorCNA ^32^ event call: green, loss; blue, neutral; red, gain. **(c)** Receiver operating characteristic (ROC) curve for kidney cancer versus healthy controls. The frozen classifier reached AUC 0.83 (95% CI, 0.51-1.00); dashed diagonal indicates chance. **(d)** Precision-recall curve for the same zero-shot evaluation. Average precision was 0.87 (95% CI, 0.60-1.00); dashed line indicates class prevalence. **(e)** AUC for LEAF-1 and a binned-coverage baseline (fragment counts over 100 kbp genomic bins) on kidney cancer detection. Error bars indicate 95% confidence intervals; dashed line denotes chance performance (AUC = 0.5).

We randomly selected 100,000 fragments per sample to be embedded by the frozen layer-4 of the LEAF-1 encoder, matching the protocol used in the previous section (**Figure 6a** and **Supplementary Figure 14**). The frozen model separated kidney cancer patients from healthy controls with AUC of 0.833 (95% CI 0.51-1.00) and average precision 0.87 (95% CI 0.60-1.00; **Figure 6c**,**d**). For comparison, a baseline classifier using a conventional binned-coverage representation (fragment counts over 100-kbp genomic bins, in place of the LEAF-1 embeddings) achieved an AUC of 0.639 (95% CI 0.31–0.89; **Figure 6e**) with the confidence interval crossing random classifier value of 0.5. Thus, a pan-cancer LEAF-1+MIL classifier trained on retrospective non-kidney cancer cohorts can detect kidney cancer signals in an independently profiled plasma cohort without retraining.

## 3. Discussion

Our results establish LEAF-1 as a fragment-level foundation model for semantic learning over DNA fragments. Instead of collapsing fragments into coordinate-bin counts, LEAF-1 represents each molecule by its sequence, assay modality, flanking context, and cleavage boundaries. A cell or sample is therefore modeled as a collection of fragment embeddings rather than as a sparse vector of genomic intervals. This preserves the fragment as the primitive unit of measurement while keeping downstream classification and pooling strategies familiar. We show that this representation approach outperforms the current leading fragmentomics-based approaches for cancer detection even on sparse datasets, establishing fragment-level, sequence-aware modeling as a strong framework for learning from cfDNA and chromatin accessibility data.

Importantly, we find that the same semantic fragment space supports different biological signals in different fragmentation chemistries. Specifically, in Tn5-based scATAC-seq, LEAF-1 embeddings preserved regulatory identity, and fragment-importance analysis linked high-scoring molecules to candidate cis-regulatory elements and transcription-factor programs. Furthermore, token and window analyses localized most scATAC signals to the fragment body, consistent with accessible regulatory sequence being the primary informative substrate. In cfDNA, by contrast, discriminative signal concentrated at the [CUT] boundary tokens, consistent with endogenous nuclease cleavage preferences and fragment-end biology driving cell type specificity. These observations suggest that LEAF-1 learns fragment features that are differentially applicable to different biological contexts and desired tasks.

This distinction is particularly important for cfDNA analysis, where fragment length, coverage, end motifs, and related aggregate features carry cancer signals. While prior fragmentomic methods often reduce each sample to engineered summaries (as discussed in the Introduction), LEAF-1 learns information-rich fragment embeddings that can directly be used in downstream classification tasks. Our evaluations under matched protocols against coordinate-binning, DELFI-style, k-mer-based, and general-purpose DNA language-model baselines strongly suggest that fragment-specific pretraining and explicit cleavage-boundary modeling capture information that is not recovered by scale alone under this task.

We further show that, beyond improved representation of individual fragments, LEAF-1 enables development of downstream models, based on multiple-instance learning (MIL), for sample-level prediction. This MIL framework becomes particularly important in the context of cfDNA-based cancer detection, where informative molecules originating from cancer cells are unlabeled and diluted within a large background of fragments from non-cancer cells. Attention-based pooling over LEAF-1 embeddings allows the classifier to weight fragments unequally while keeping the encoder frozen, which, as we showed, results in increased sensitivity and specificity at extremely shallow genome coverages. The resulting pan-cancer performance and fragment-efficiency gains support the premise that modeling cfDNA as a distribution of molecules is more informative than treating it only as coordinate coverage.

Our independently profiled kidney cohort provides an initial test of zero-shot transfer beyond the supervised training cancer types. To our knowledge, this is the first demonstration that a cfDNA model trained on one set of cancer types can detect an unseen cancer type in an independently profiled cohort without retraining. This result should at the same time be interpreted cautiously: our retrospective cohort was small, and the resulting confidence intervals were wide. Nonetheless, even at the lower end of the observed confidence intervals, our finding supports the notion that LEAF-1 captures generalizable cancer-associated fragmentation signals that can transfer across malignancies. By enabling reliable detection from low-input, low-depth cfDNA, LEAF-1 could reduce assay cost and turnaround time, broaden access to liquid-biopsy testing, and extend its utility to early-stage and low-shedding cancers, settings in which current ctDNA assays remain limited.

Future versions of LEAF-1 could expand both model scale and assay coverage. Specifically, we envision that incorporating additional fragmentation chemistries would broaden the scope of semantic fragment learning to other DNA fragment-based assays. In cfDNA, for example, incorporating fragment-level methylation information may further improve detection and tissue-of-origin inference. More broadly, our results support semantic learning over individual DNA fragments as a way to preserve biochemical, cell-associated, and disease-associated signals that are otherwise lost during coordinate-based aggregation.

## 4. Methods

### 4.1. Datasets and pretraining corpus

The pretraining corpus spanned three DNA-fragmentation modalities: cell-free DNA, bulk ATAC-seq, and single-cell ATAC-seq. Across modalities, the corpus contained 2,620 unique samples, 601,093 single cells, and 58.1 billion sequenced fragments before modality balancing. The cfDNA component contained 2,125 plasma samples across several studies: Adalsteinsson et al. (^32^; 1,357 samples), Cristiano et al. (^5^; 538 samples), Jiang et al. (^33^; 202 samples), and Sun et al. (^34^; 28 samples), totaling 31.4 billion fragments. The bulk ATAC-seq component contained 404 TCGA samples from Corces et al. ^19^ totaling 22.4 billion fragments and was used for pretraining only in this manuscript. The scATAC component came from the single-cell atlas of Zhang et al. ^20^, including GTEx, LungMap, and ENCODE components, and contained 91 tissue samples, 601,093 cells, and 4.3 billion fragments across 108 cell types.

To balance exposure across modalities during pretraining, samples were replicated to match the cfDNA sample count in each modality (**Supplementary Table 1**). File order was shuffled between training iterations, and file-level sampling used seven fragments per file to diversify exposure across samples and modalities. A subset of single-cell samples was held out for pretraining validation. Downstream evaluations used unique samples or cells without modality-balancing replication and were analyzed under task-specific cross-validation protocols.

The main cfDNA benchmark used the Cristiano cohort of 538 samples from seven cancer types plus healthy controls. The scATAC downstream analysis used 108 cell types across 28 tissues; for balanced cell-level evaluation, up to 100 cells per cell type were retained where available, yielding 10,132 cells with cell-type counts ranging from 67 to 100. The independent kidney cohort contained 12 cfDNA samples, six healthy and six kidney cancer, collected and processed independently of the retrospective cohorts used for pretraining and downstream Cristiano training.

### 4.2. Fragment preprocessing and tokenization

All genomic coordinates were referenced to GRCh38. Fragment files were filtered to retain alignments with mapping quality at least 30 and fragment length at least 20 bp. Fragments overlapping ENCODE blacklist regions were excluded to reduce artifacts from repetitive or poorly mappable loci. Each fragment was represented by left flanking sequence, the fragment body, two explicit [CUT] boundary tokens, and right flanking sequence. Inputs used a 2,048-token budget, with flank lengths dynamically adjusted to fill the available budget while retaining at least 100 tokens per side when possible. Fragments near chromosome boundaries were handled by maximally extending the accessible flank and compensating with the opposite flank when possible. Strand orientation was randomized with 50% probability during pretraining.

Tokenization used a byte-pair encoding vocabulary trained on a representative genomic corpus spanning all three modalities. The tokenizer used a 4,096-token vocabulary plus special tokens for sequence boundaries, masking, padding, unknown sequence, cleavage boundaries, and modality identifiers. Each input followed the structure [CLS] [MOD] left_flank [CUT] body [CUT] right_flank. The [MOD] token identified the fragmentation modality, allowing a shared encoder to learn modality-conditioned patterns across endogenous-nuclease cfDNA and Tn5-based accessibility data.

### 4.3. Model architecture and pretraining

LEAF-1 is a 12-layer transformer encoder with 12 attention heads per layer, 64 dimensions per head, hidden dimension 768, and feedforward dimension 3,072 with GELU activations. The model contains approximately 90 million parameters. Positional embeddings were learned, and modality information was supplied through explicit modality tokens rather than token-type embeddings.

Pretraining used masked language modeling. Base tokens were masked with probability 0.15, while [CUT] tokens were masked with probability 0.30 to emphasize cleavage-boundary prediction. Both [CUT] tokens flanking a fragment were masked jointly so that the model could not infer one boundary trivially from the other. For masked positions, 80% were replaced with [MASK], 10% with random vocabulary tokens, and 10% were left unchanged. Special tokens other than [CUT] were not masked. The training loss was cross-entropy over masked positions only.

Training proceeded for 40,000 steps with effective batch size 4,096 sequences, using 32 sequences per GPU, 32 gradient-accumulation steps, and 4 GPUs when available. This corresponded to approximately 8.4 million tokens per update and 163.8 million sampled fragments processed during training. Optimization used AdamW with learning rate 1e-4, linear warmup over 1,000 steps, linear decay to zero, weight decay 1e-5, mixed precision, and gradient clipping at maximum norm 1.0. Training continued for the full schedule without early stopping. Random seeds were fixed at 42 for Python, NumPy, PyTorch, and scikit-learn operations.

### 4.4. Multi-window embedding extraction

Embeddings were extracted from the frozen encoder by partitioning each tokenized fragment into left flank, fragment body, and right flank according to the [CUT] token positions. Within each region, adaptive average pooling divided the variable-length token sequence into three positional windows, yielding nine regional descriptors. These were concatenated with four special-token embeddings, [CLS], [MOD], [CUT1], and [CUT2]. With hidden dimension 768, the resulting fragment representation had dimension 9,984: nine pooled windows plus four special-token embeddings.

Embeddings were extracted from the embedding layer and all 12 transformer layers during layer-selection experiments. Support-vector-machine evaluations across downstream tasks identified layer 4 as the primary downstream layer, balancing local sequence information from early layers with contextual information from deeper layers. The main analyses therefore used frozen layer-4 embeddings. cfDNA analyses sampled 100,000 fragments per sample unless otherwise specified; scATAC analyses used up to 1,000 fragments per cell to reflect the sparsity of single-cell accessibility data.

### 4.5. Cross-validation and leakage control

Downstream evaluations used nested cross-validation unless a frozen external validation model was being evaluated. The outer loop was reserved for performance estimation, and the inner loop was used for hyper-parameter selection. Feature normalization parameters were estimated only from training data within each fold and then applied to validation or test data. After hyperparameter selection, models were fit using the corresponding training data and evaluated on held-out outer folds or independent samples.

For cfDNA representation benchmarks, matched leave-one-out nested cross-validation was used across LEAF-1, coordinate-binning, k-mer, DELFI-style, and external DNA-language-model baselines. For scATAC classification, 10-fold stratified cross-validation was used with inner 5-fold hyperparameter selection. Sex-specific comparisons excluded inappropriate controls or comparison samples within the fold construction where needed, preventing classifiers from exploiting sex-chromosome dosage rather than disease signal.

### 4.6. scATAC downstream analysis

For cell-level scATAC prediction, each cell was represented by the mean of its frozen LEAF-1 fragment embeddings. One-vs-all classifiers were trained for 108 cell types, with logistic regression used for the main analysis and linear support-vector-machine models used in matched supplementary comparisons where appropriate. Coordinate-binning baselines were evaluated under matched folds so that differences reflected representation rather than split composition. Hyperparameters were selected within inner folds by validation AUC.

For candidate cis-regulatory-element analysis, fragment-level classifier scores were summarized at case-associated candidate cCREs for each cell type and compared with corresponding scores in control cell types. The main paired comparison used a two-sided paired t-test to assess whether case-associated cCREs had higher mean fragment scores than controls. For motif/accessibility concordance, we performed a three-step analysis. First, for each of 102 one versus all cell type comparisons, we evaluated the association between transcription factor (TF) motif presence and the complete distribution of LEAF-1 fragment scores. Using FIRE ^21^, we computed a motif enrichment statistic based on mutual information between motif occurrence and fragment score, generating a cell-type-specific enrichment value for each TF. Positive values indicate preferential occurrence of a motif in highly scored fragments, whereas negative values indicate enrichment among low-scoring fragments. Second, independent of the model, we estimated TF promoter accessibility from the raw scATAC-seq data. Tn5 insertions within promoter regions were aggregated across cells of the same type to generate pseudobulk profiles, normalized to counts per million, and log-transformed, yielding a promoter accessibility measure for each TF across cell types. Third, for each TF, we calculated the Pearson correlation between motif enrichment scores and promoter accessibility across the 102 cell-type comparisons. Significant TF motif associations across cell-type comparisons were summarized together with accessibility direction, allowing motifs to be grouped into positive and negative model-score programs and compared with chromatin-accessibility changes.

### 4.7. cf DNA representation benchmarks

For cf DNA representation benchmarks, frozen LEAF-1 embeddings were evaluated under the same matched leave-one-out nested cross-validation protocol as all baselines. Linear classifiers were trained on mean-pooled sample-level LEAF-1 embeddings for the representation benchmark panels, separating the quality of the frozen representation from the later MIL pooling model. The 14 cfDNA detection tasks included cancer-versus-healthy and cancer-type comparisons derived from the Cristiano cohort.

Coordinate-binning baselines included Bin-only features at 575, 1K, 3K, and 10K bins; Bin+Length features using a 150 bp short/long fragment-length stratification; DELFI-style fragmentation features; and Bin+DELFI feature combinations. Canonical k-mer frequency vectors were reverse-complement folded and evaluated for k = 3 to 8, with k = 3 to 6 emphasized in the main figure and the full k-mer table retained in the supplement. External DNA language model baselines included DNABERT-2, DNABERT-S, GENA-LM, NT-v2, and Evo 2 and were evaluated with the same fold assignments and model-selection protocol.

### 4.8. LEAF-1+MIL cf DNA model

For sample-level cfDNA cancer detection, each sample was treated as a bag of 100,000 frozen LEAF-1 fragment embeddings. The MIL model consisted of an optional linear feature projector with layer normalization, a gated-attention pooling module, and an elastic-net classification head. The feature projector mapped the 9,984-dimensional fragment embeddings to 512 or 1,024 dimensions. The gated-attention module computed instance weights from the element-wise product of tanh and sigmoid branches, followed by softmax normalization over fragments in the bag.

Attention pooling over very large fragment bags can reduce the variance of the pooled representation, because weighted averages over many correlated fragment embeddings tend toward their shared component. To avoid this variance collapse, the pooled representation was rescaled using the Herfindahl-Hirschman effective sample size. For attention weights w_i, n_eff = 1 / sum_i w_i^2. If h is the unscaled weighted mean of projected fragment embeddings, the rescaled representation was h’ = h * sqrt(n_eff). This leaves sharply concentrated attention close to unscaled while restoring dynamic range when attention is distributed over many fragments.

The MIL classification head used elastic-net regularization with C in {1, 3.16, 10, 31.6, 100} and alpha in {0.0, 0.3}. Projector and attention regularization used either dropout rates in {0.0, 0.1, 0.2} or ridge penalties with C in {0.316, 1, 3.16, 10}, but not both simultaneously. The reduced hyperparameter grid contained 80 configurations per fold. MIL training used Adam with linear warmup from 1e-6 to 5e-4 over 5 epochs, a 15-epoch plateau, and cosine decay to 1e-6 up to 200 epochs. Class imbalance was handled with weighted random sampling. Early stopping monitored an exponential moving average of validation AUC with patience 20 epochs and minimum improvement 0.005, and predictions were averaged from the top three validation checkpoints.

### 4.9. Single-component and subsampling analyses

Layer analyses evaluated embeddings from the embedding layer and all 12 transformer layers using matched downstream classifiers. Segment and window analyses trained classifiers on restricted feature subsets corresponding to left flank, fragment body, right flank, [CLS], [MOD], [CUT1], [CUT2], and the nine individual pooled windows. These analyses were run within each modality-specific benchmark to determine whether discriminative signal was concentrated in fragment bodies, flanks, or cleavage-boundary tokens.

Fragment-count subsampling evaluated cfDNA performance as a function of the number of fragments available per sample, varying from 1,000 to 100,000 fragments. LEAF-1+MIL and coordinate-binning baselines were evaluated under matched fold assignments and fragment-count settings. Fragment-efficiency gains were computed per cancer type as the fold reduction in fragment count required by LEAF-1 embeddings to match the coordinate-binning AUC at 100,000 fragments.

### 4.10. Stratification of detection performance by ctDNA level

To assess how classification performance varies with circulating tumor DNA (ctDNA) burden, we used ichorCNA estimated tumor fractions as a proxy for ctDNA burden. These tumor fraction estimates were obtained from Doebley et al. ^31^, who reanalyzed the DELFI cohort originally described by Cristiano et al. ^5^. For each sample, we obtained model predicted scores from LEAF-1 and DELFI. DELFI was represented by a logistic regression classifier trained on DELFI features extracted from the same 100k-fragment input. Among cancer samples with an available matched tumor fraction estimate, we defined three ctDNA-burden groups, low, medium, and high, using cohort-wide tertiles of ichorCNA tumor fraction. We first compared the distributions of LEAF-1 predicted positive probabilities across ctDNA-burden groups to assess whether model confidence increased with inferred ctDNA burden. We then evaluated classification performance within each ctDNA-burden group. For each of the seven cancer types included in this analysis, bile duct, breast, colorectal, gastric, lung, ovarian, and pancreatic cancer, and within each ctDNA-burden group, we computed a cancer-versus-healthy AUC by comparing cancer samples of that cancer type in the given ctDNA-burden group with the full set of healthy controls. This procedure yielded one paired LEAF-1 and DELFI AUC per cancer type within each ctDNA-burden group. To test whether LEAF-1 outperformed DELFI within each ctDNA-burden group, we applied a two-sided paired Wilcoxon signed-rank test to the seven paired per-cancer AUC differences.

### 4.11. Independent kidney validation protocol

For the independent kidney cohort, plasma was collected and processed outside the retrospective cohorts. Plasma samples for 6 RCC patients were provided by the McGill RCC biobank, while plasma samples for 6 healthy controls were collected from volunteers recruited for this study. All samples were received following obtaining written consents from the participants and after approval of the study by the McGill University Health Centre Research Ethics Board (MUHC REB # 2015-1257). 9mL of blood was collected in a K2 EDTA tube, inverted 10 times then kept at 4°C. Plasma was isolated by centrifugation (2000xg, 15min, 4°C) within 1 hour of collection, aliquoted in cryovials and stored at -80°C. cfDNA was extracted from 2-4mL of re-centrifuged (2000xg, 10min) plasma using the Maxwell RSC Rapid ccfDNA Kit (Promega). The quantity and quality of the cfDNA were measured by Qubit (Life Technologies) and Tapestation cf DNA ScreenTape assay (Agilent) respectively.

cfDNA libraries were prepared from 10 ng input using the Watchmaker DNA Library Prep Kit for Fragmented Double-Stranded DNA and sequenced on an Illumina MiSeq v2 with paired-end 150 bp reads. To assess copy-number profiles and estimate tumor fraction, ichorCNA ^32^ was applied to all samples. For the LEAF-1 analysis,100,000 fragments were randomly sampled per sample with seed 42, embedded with the frozen layer-4 encoder, and evaluated with frozen Cristiano-trained MIL models. Predictions were averaged across fold models, and no model parameters, thresholds, or preprocessing choices were adjusted after receiving the independent samples. As a baseline, we constructed a fragment-count feature matrix over the same samples using 100 kbp genomic bins and applied a frozen Cristiano-trained binning model for comparison with LEAF-1.

### 4.12. Performance metrics and statistics

Primary metrics were area under the receiver operating characteristic curve (AUC-ROC) and average precision. Average-precision lift was computed relative to class prevalence. Bootstrap confidence intervals used bias-corrected and accelerated intervals for paired representation benchmarks and 10,000-resample bootstrap intervals for primary clinical metrics unless otherwise specified. Paired bootstrap deltas summarized LEAF-1 improvements over baselines using matched test predictions. Group separation in the independent kidney cohort was assessed with the Mann-Whitney U test and Cohen’s d effect size.

## Data and code availability

Processed data for the in-house cell-free DNA (cfDNA) samples and associated sample metadata for the ccRCC and healthy control samples have been deposited in Zenodo and are available at https://doi.org/10.5281/zenodo.21269430. LEAF-1 is available at https://github.com/csglab/leaf-1. The code for reproducing the analyses shown in the manuscript is available at https://github.com/csglab/leaf-1-figures.

## 5. Acknowledgements

We thank Chiara Ricci-Tam for aiding with visualization and figure preparation, and April Pawluk for providing feedback on the manuscript draft. This work was supported by grants from the Canadian Institutes of Health Research (CIHR) [PJT-173317] to HSN, research support provided by Arc Institute to HG, and resources provided by Calcul Québec (calculquebec.ca) and the Digital Research Alliance of Canada (alliancecan.ca) to HSN. HSN holds a CIHR Canada Research Chair. HG is a Core Investigator at Arc Institute.

## 6. Author contributions

HH: Conceptualization, Methodology, Software, Validation, Formal analysis, Investigation, Data Curation, Writing - Original Draft, Writing - Review & Editing, Visualization; JZ: Methodology, Validation, Formal analysis, Investigation, Data Curation, Writing - Original Draft, Writing - Review & Editing, Visualization; MA: Methodology, Investigation, Writing - Review & Editing; LY: Investigation; ST: Resources, Writing - Review & Editing; YR: Investigation, Resources, Writing - Review & Editing; HG: Conceptualization, Methodology, Investigation, Resources, Writing - Original Draft, Writing - Review & Editing, Supervision; HSN: Conceptualization, Methodology, Investigation, Resources, Writing - Original Draft, Writing - Review & Editing, Supervision, Funding acquisition.

