## Supplementary Information for "Semantic fragment representations for coordinate-free analysis of genomics data"

Hamed Heydari[1,2,3,#], Jing Zhao[1,2,#], Madeleine Arseneault[2], Leila Younesian[2], Simon Tanguay[4], Yasser Riazalhosseini[1,2], Hani Goodarzi[3,5,6,7,8,*], Hamed Najafabadi[1,2,9,10,*]

[1]Department of Human Genetics, McGill University, Montreal, QC, H3A 0C7, Canada
[2]Victor P. Dahdaleh Institute of Genomic Medicine, Montreal, QC, H3A 0G1, Canada
[3]Arc Institute, Palo Alto, CA, US
[4]Division of Urology, McGill University, Montreal, Quebec, Canada
[5]Department of Biochemistry and Biophysics, University of California, San Francisco, CA, US
[6]Department of Urology, University of California, San Francisco, San Francisco, California, USA
[7]Helen Diller Family Comprehensive Cancer Center, University of California, San Francisco, CA, US
[8]Bakar Computational Health Sciences Institute, University of California, San Francisco, CA, US
[9]McGill Centre for RNA Sciences, McGill University, Montreal, QC, Canada
[10]Goodman Cancer Institute, McGill University, Montreal, QC, Canada

[#]These authors contributed equally to this work.

### Supplementary tables

**Supplementary Table 1.** Composition of training and evaluation datasets across three modalities.

| Modality | Source | Studies / Types | Samples | Cells | Total Fragments |
|---|---|---|---|---|---|
| Cell-free DNA | Adalsteinsson et al., 2017[1] | 2 | 1,357 | – | 20.1B |
| | Cristiano et al., 2019[2] | 7 | 538 | – | 7.9B |
| | Jiang et al., 2015[3] | 4 | 202 | – | 3.0B |
| | Sun et al., 2019[4] | 2 | 28 | – | 0.4B |
| | *Subtotal* | *15* | *2,125* | – | *31.4B* |
| Bulk ATAC-seq | TCGA[5] | 23 | 404 | – | 22.4B |
| Single-cell ATAC | Zhang et al., 2021[6] | 28 | 91 | 601,093 | 4.3B |
| **Total** | – | **66** | **2,620** | **601,093** | **58.1B** |

**Supplementary Table 2.** Clinical and sequencing characteristics of the independent ccRCC validation cohort.

| Group | Sample ID | Sex | Pathologic Diagnosis | pT Stage | pN Stage | pM Stage | Nuclear Grade | First Metast. Site | Disease Status | Tissue Specimen Source / Procedure | cfDNA Frag. Count |
|---|---|---|---|---|---|---|---|---|---|---|---|
| Healthy Control | HC001 | Female | N/A | – | – | – | – | – | Healthy | – | 875616 |
| | HC007 | Male | N/A | – | – | – | – | – | Healthy | – | 961478 |
| | HC009 | Male | N/A | – | – | – | – | – | Healthy | – | 926346 |
| | HC010 | Female | N/A | – | – | – | – | – | Healthy | – | 865937 |
| | HC012 | Female | N/A | – | – | – | – | – | Healthy | – | 937695 |
| | HC013 | Male | N/A | – | – | – | – | – | Healthy | – | 1190313 |
| ccRCC Patient | PID038 | Male | Clear Cell (Conventional) Renal Cell Carcinoma | pT3a | pNx | NA | 3/4 | None | NED | Right kidney radical nephrectomy | 882795 |
| | PID039 | Male | Clear Cell (Conventional) Renal Cell Carcinoma | pT3a | pNx | NA | 3/4 | None | NED | Right kidney partial nephrectomy | 980599 |
| | PID019 | Male | Clear Cell (Conventional) Renal Cell Carcinoma | pT3a | pN0 | Mx | 3/4 | Brain | NED | Right kidney radical nephrectomy | 1371154 |
| | PID040 | Male | Clear Cell (Conventional) Renal Cell Carcinoma | pT3a | pNx | Mx | 3/4 | None | NED | Right kidney radical nephrectomy | 989556 |
| | PID020 | Male | Clear Cell (Conventional) Renal Cell Carcinoma with extensive necrosis | pT3a | pNx | M1 | 3/4 | Lung (2017) | AWD | Right kidney radical nephrectomy | 1219710 |
| | PID041 | Female | Clear Cell (Conventional) Renal Cell Carcinoma | pT3a | pNx | M0 | 4/4 | None | NED | Right kidney nephrectomy | 1610306 |

### Supplementary figures

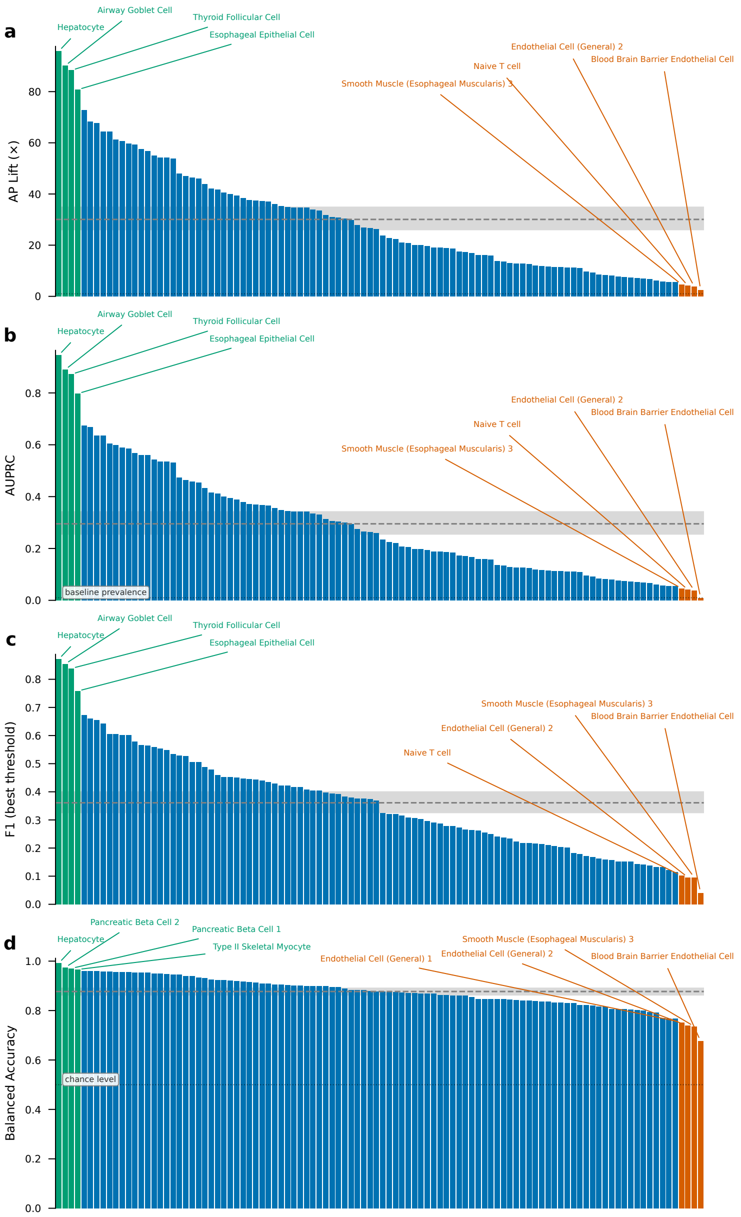


**Supplementary Figure 1.** Per-cell-type scATAC classification performance across complementary metrics. One-vs-all classification performance for 108 cell types across 28 tissues, using mean-pooled LEAF-1 embeddings from the frozen encoder with up to 1,000 fragments per cell[6] (logistic regression; see **Methods**). Each bar is one cell type, ranked from highest to lowest within each panel; green bars indicate top-performing cell types and orange bars indicate the lowest-performing or broadly defined cell types, with representative examples labeled. **(a)** Average-precision lift over class prevalence (fold). **(b)** Area under the precision-recall curve (AUPRC); solid line marks baseline prevalence. **(c)** F1 score at the best threshold. **(d)** Balanced accuracy; solid line marks chance level. In all panels, the dashed line indicates the across-cell-type mean and the shaded band the interquartile range.

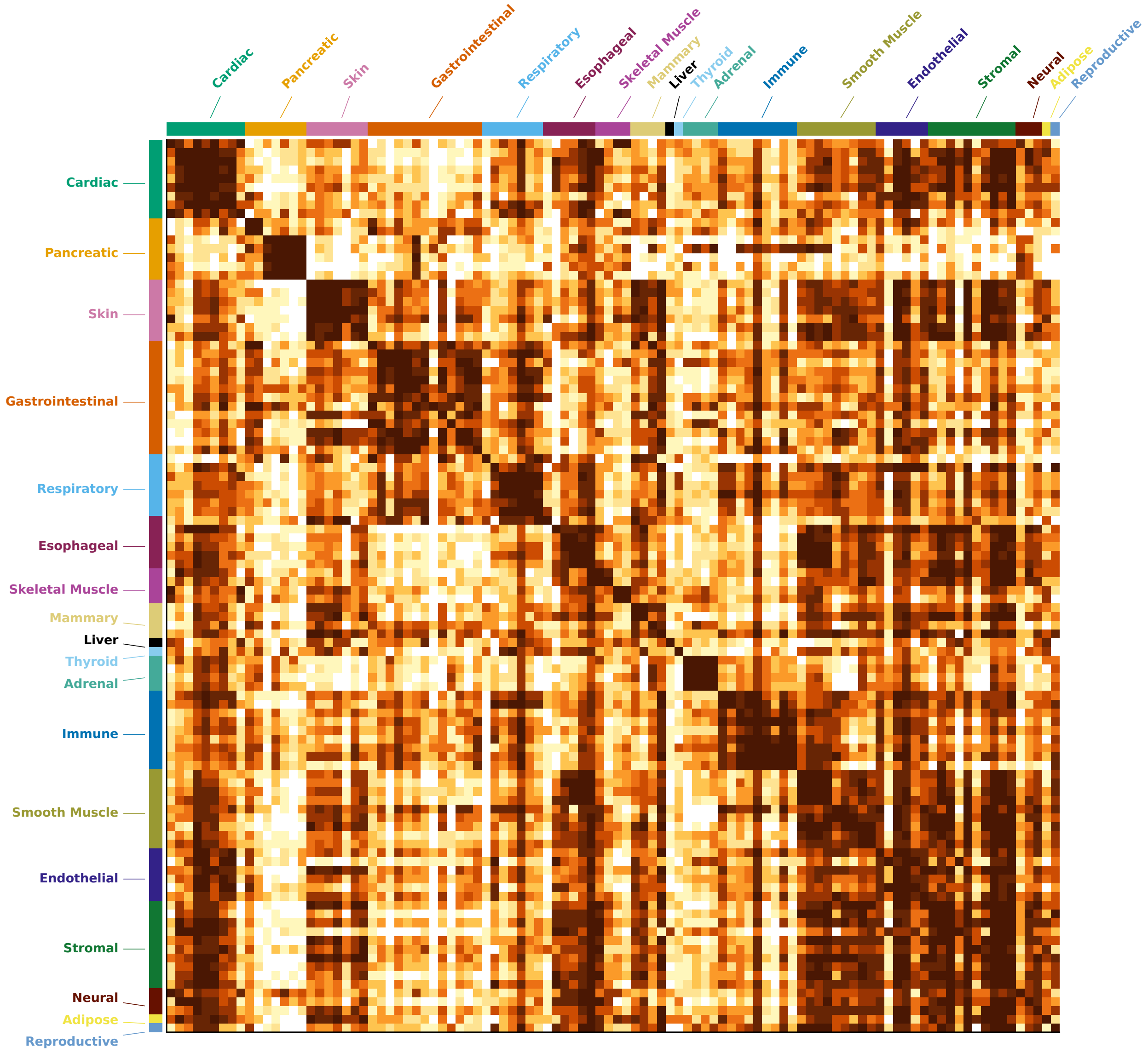


**Supplementary Figure 2.** Confusion matrix for scATAC cell-type classification. Confusion matrix for one-vs-all classification of 108 cell types across 28 tissues using mean-pooled LEAF-1 embeddings from the frozen encoder with up to 1,000 fragments per cell[6] (see **Methods**). Rows and columns correspond to true and predicted cell types, ordered by tissue category; the color bars along the top and left annotate tissue category (Cardiac, Pancreatic, Skin, Gastrointestinal, Respiratory, Esophageal, Skeletal Muscle, Mammary, Liver, Thyroid, Adrenal, Immune, Smooth Muscle, Endothelial, Stromal, Neural, Adipose, and Reproductive). Color intensity encodes classification frequency from low (light) to high (dark), with the diagonal corresponding to agreement between true and predicted labels.

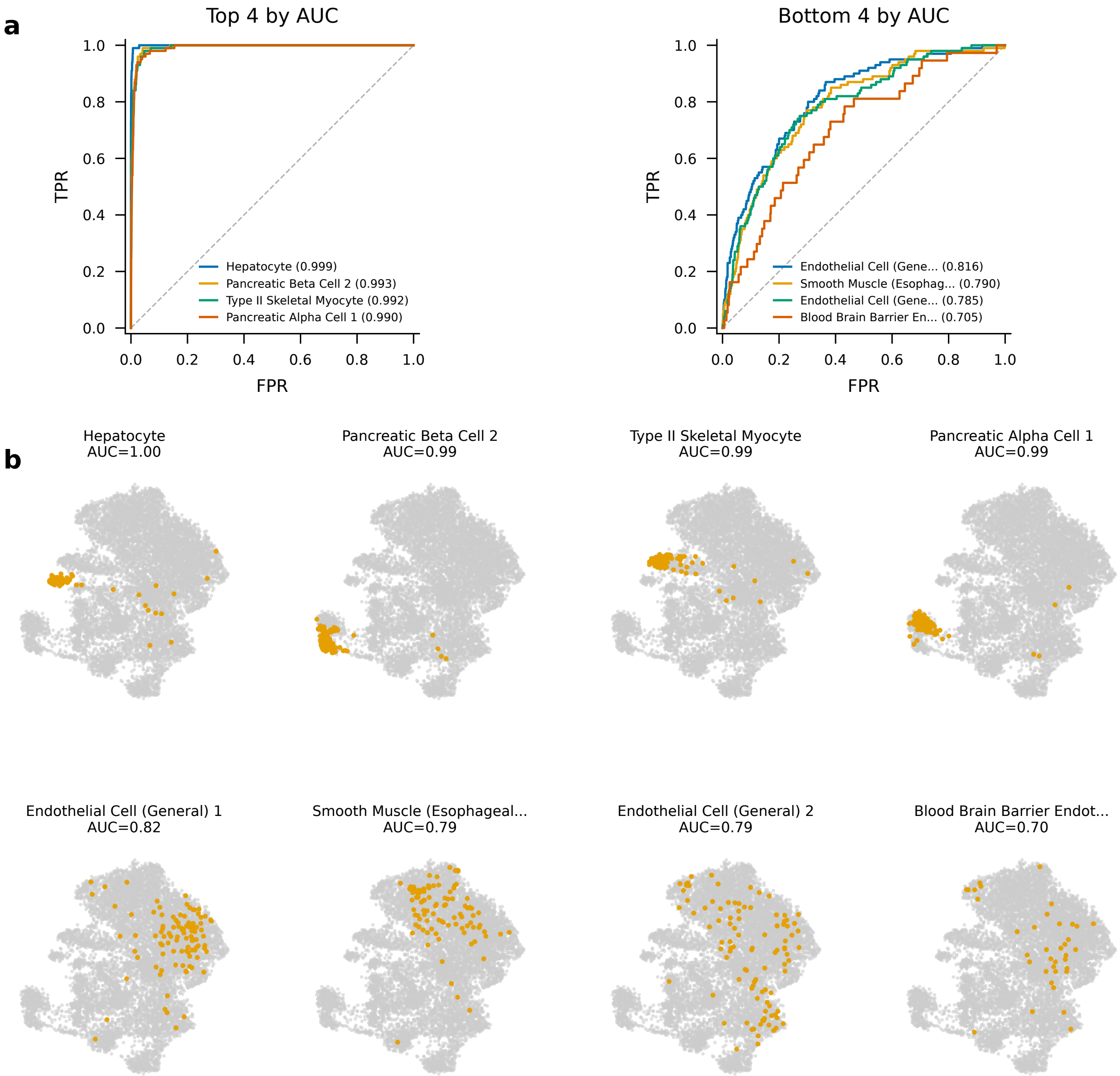


**Supplementary Figure 3.** ROC curves and embedding localization for the best- and worst-classified scATAC cell types. Per-cell-type performance for one-vs-all classification of scATAC cell types using mean-pooled LEAF-1 embeddings from the frozen encoder with up to 1,000 fragments per cell[6] (see **Methods**). **(a)** Receiver-operating-characteristic curves for the four highest-AUC cell types (left: hepatocyte, 0.999; pancreatic beta cell 2, 0.993; type II skeletal myocyte, 0.992; pancreatic alpha cell 1, 0.990) and the four lowest-AUC cell types (right: endothelial cell (general) 1, 0.816; smooth muscle (esophageal muscularis) 3, 0.790; endothelial cell (general) 2, 0.785; blood-brain-barrier endothelial cell, 0.705). Dashed diagonal indicates chance. **(b)** UMAP projections of mean-pooled cell embeddings for the same eight cell types; in each panel, cells of the indicated cell type are highlighted in orange against all other cells in grey, with the corresponding AUC shown above each plot.

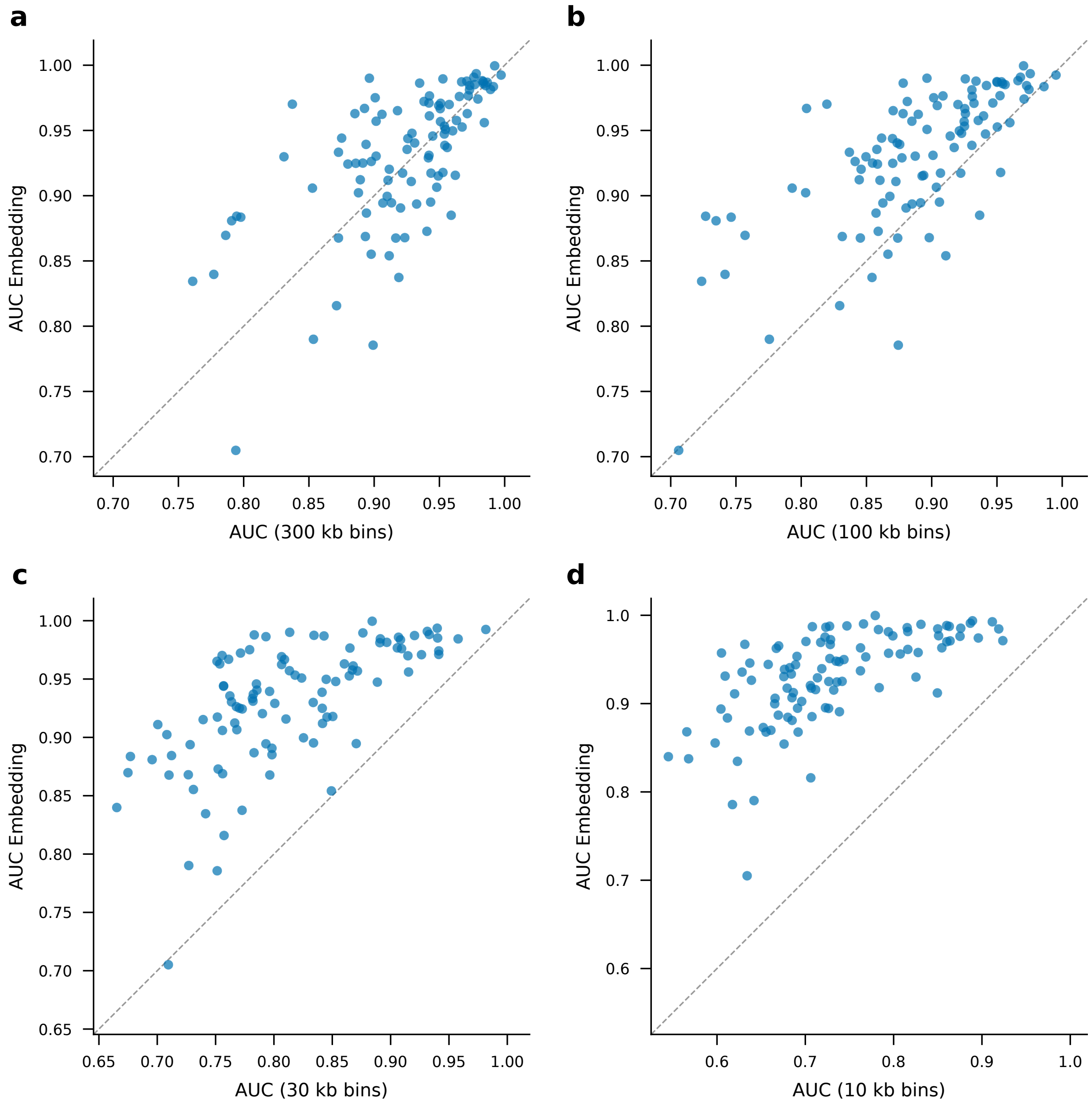


**Supplementary Figure 4.** LEAF-1 embeddings versus coordinate-binning across bin resolutions in single-cell ATAC-seq. Per-cell-type one-vs-all classification AUC for mean-pooled LEAF-1 embeddings (y-axis) versus coordinate-binning features (x-axis), evaluated under matched cross-validation with up to 1,000 fragments per cell[6] (see **Methods**). Each point is one cell type. **(a)** 300kb bins. **(b)** 100kb bins. **(c)** 30kb bins. **(d)** 10kb bins. The dashed diagonal indicates equal performance between the two representations.

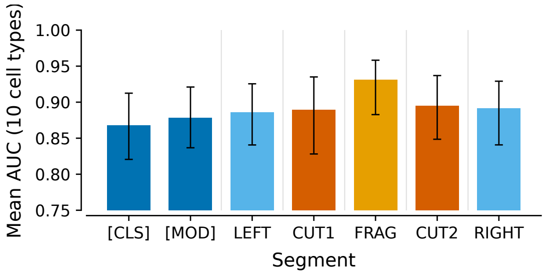


**Supplementary Figure 5.** Single-segment analysis of LEAF-1 token components in single-cell ATAC-seq. Mean one-vs-all classification AUC across 10 cell types for classifiers trained on individual LEAF-1 segment embeddings, using up to 1,000 fragments per cell[6] (see **Methods**). Bars correspond to the [CLS] token, [MOD] token, left flank (LEFT), first cleavage-boundary token (CUT1), fragment body (FRAG), second cleavage-boundary token (CUT2), and right flank (RIGHT). Error bars indicate variability across the 10 cell types.

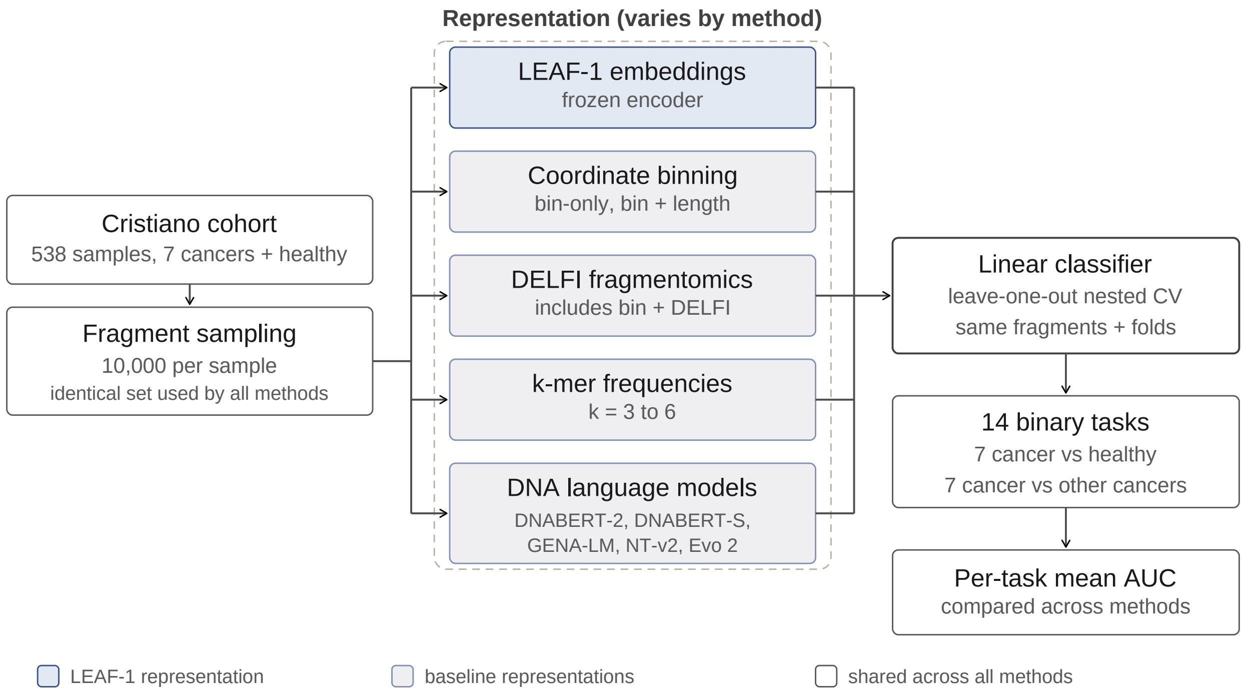


**Supplementary Figure 6.** Design of the matched cfDNA representation benchmark. Schematic of the controlled comparison used to evaluate LEAF-1 against baseline representations on cfDNA from the Cristiano cohort[2] (538 samples; seven cancer types and healthy controls). For each sample, the same 10,000 fragments were used by every method, so that arms differed only in representation. Each representation was passed to a linear classifier (logistic regression) trained under matched leave-one-out nested cross-validation, with identical fragments and fold assignments across methods (see **Methods**). The representations were LEAF-1 embeddings from the frozen encoder, coordinate binning (bin-only and bin + fragment length at multiple resolutions), DELFI-style fragmentomic features (including bin + DELFI combinations), canonical k-mer frequencies (k = 3 to 6), and five general-purpose DNA language models (DNABERT-2[7], DNABERT-S[8], GENA-LM[9], NT-v2[10] and Evo 2[11]). Classifiers were evaluated on 14 binary tasks, comprising seven cancer-versus-healthy and seven cancer-versus-other-cancers (tissue-of-origin) comparisons, summarized as per-task mean AUC. Box color indicates the LEAF-1 representation, baseline representations, and components shared across all methods, as shown in the legend.

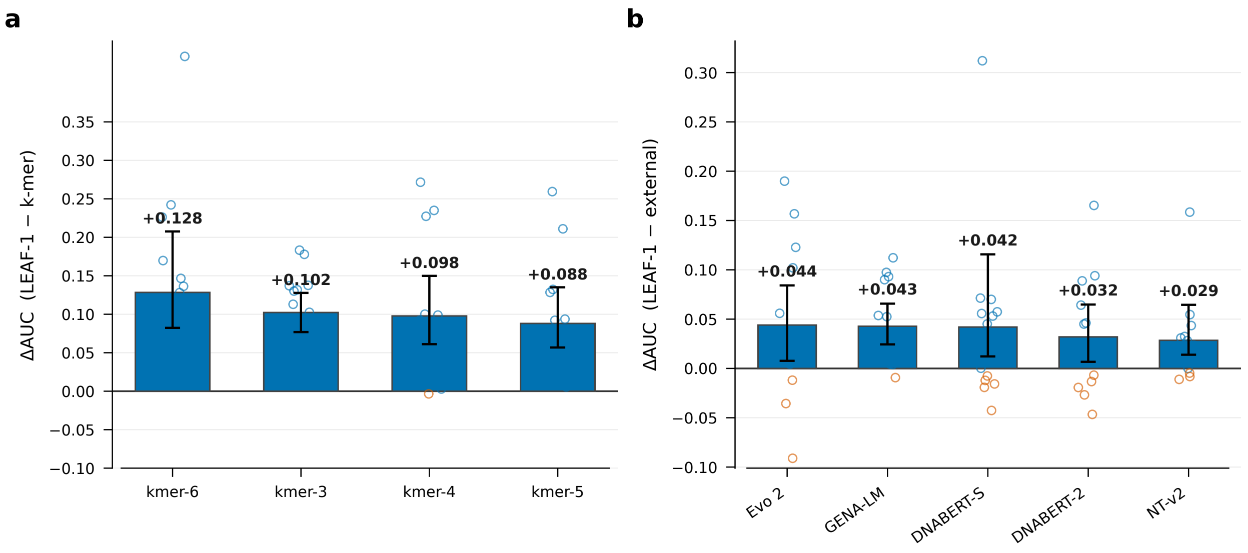


**Supplementary Figure 7.** LEAF-1 versus k-mer and DNA language-model baselines in cfDNA. Paired per-task differences in AUC (ΔAUC) between LEAF-1 embeddings and each baseline representation on the Cristiano cohort[2], evaluated under the matched protocol with identical fragments and fold assignments across methods (**Supplementary Figure 6**; see **Methods**). Each bar shows the mean ΔAUC across the 14 binary tasks (seven cancer-versus-healthy and seven cancer-versus-other-cancers); error bars indicate 95% bootstrap confidence intervals. Each point is one task. **(a)** Canonical k-mer frequency baselines (k = 3 to 6). **(b)** Five general-purpose DNA language models (Evo 2[11], GENA-LM[9], DNABERT-S[8], DNABERT-2[7], NT-v2[10]). Positive values indicate higher AUC for LEAF-1; the solid line at zero marks equal performance.

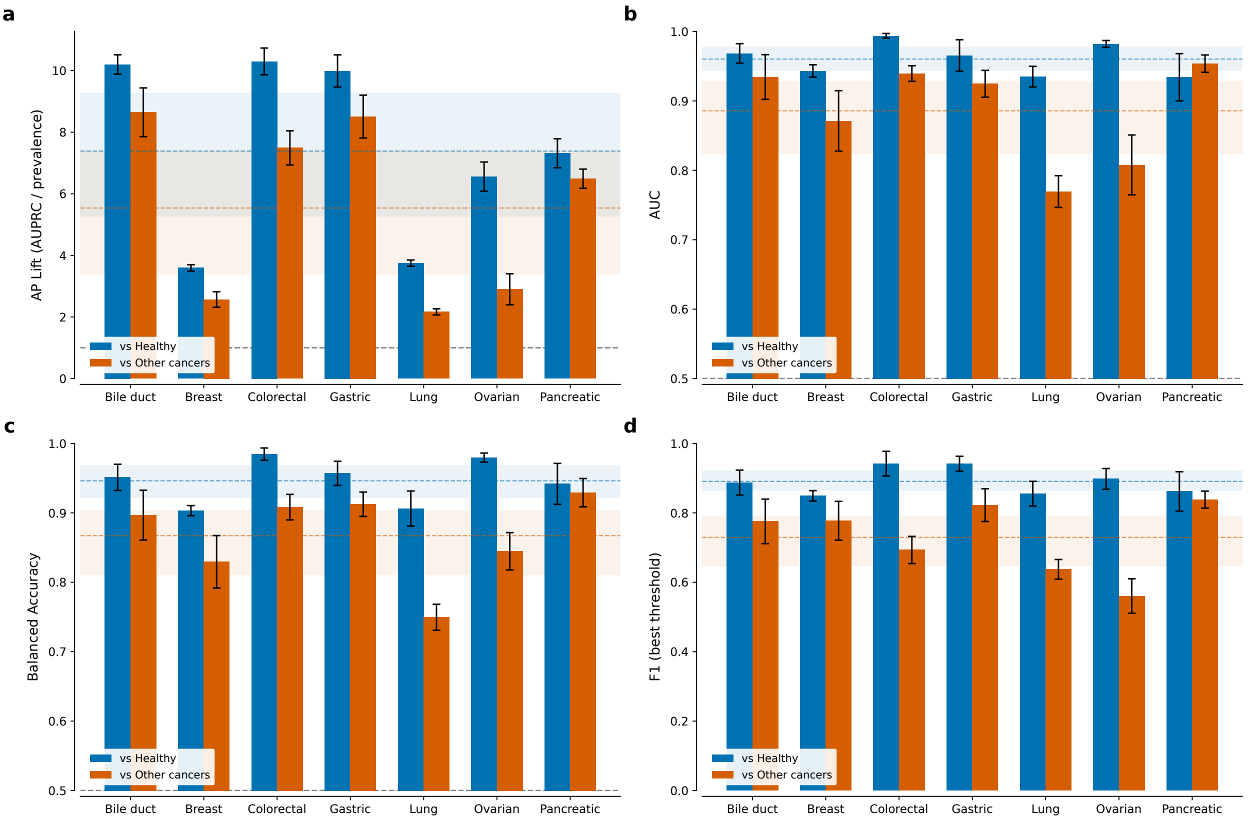


**Supplementary Figure 8.** Per-cancer cfDNA detection and discrimination across complementary metrics. LEAF-1+MIL performance for each of seven cancer types in the Cristiano cohort[2], using 100,000 fragments per sample (see **Methods**). In all panels, blue bars show cancer-versus-healthy detection and orange bars show discrimination of each cancer type from the other cancer types; error bars indicate 95% bootstrap confidence intervals; the blue and orange dashed lines indicate the across-cancer mean for the detection and discrimination tasks, respectively, with shaded bands showing the confidence interval. **(a)** Average-precision lift over class prevalence; grey dashed line marks no lift (lift = 1). **(b)** AUC; grey dashed line marks chance. **(c)** Balanced accuracy; grey dashed line marks chance. **(d)** F1 score at the best threshold.

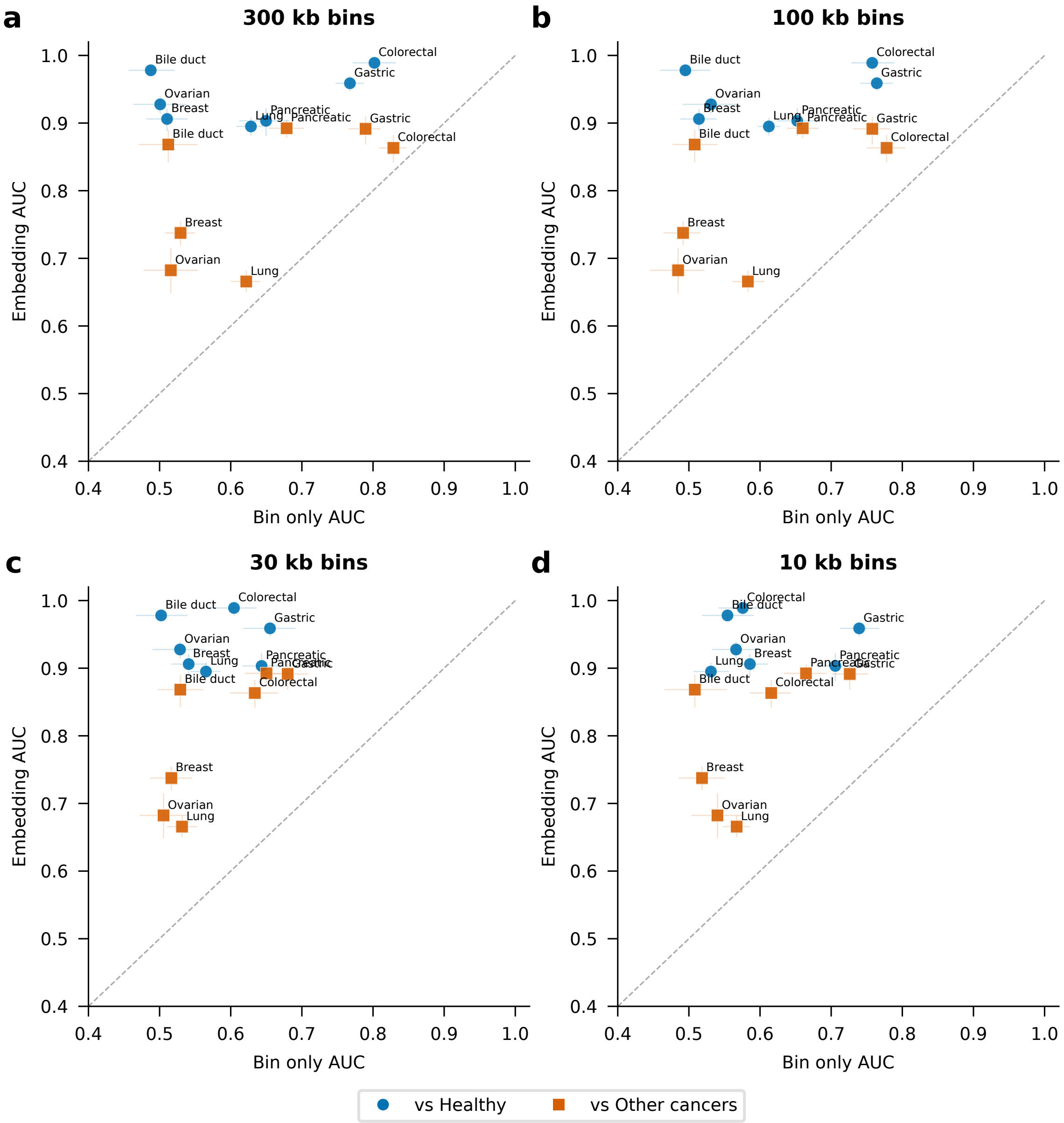


**Supplementary Figure 9.** LEAF-1+MIL embeddings versus Bin-only features across bin resolutions in cfDNA. Per-cancer AUC for LEAF-1+MIL embeddings (y-axis) versus Bin-only coordinate features (x-axis) in the Cristiano cohort[2], evaluated under matched cross-validation (see **Methods**). Each point is one cancer type; blue circles show cancer-versus-healthy detection and orange squares show cancer-versus-other-cancers discrimination. Error bars indicate confidence intervals along both axes, and the dashed diagonal indicates equal performance between the two representations. **(a)** 300kb bins. **(b)** 100kb bins. **(c)** 30kb bins. **(d)** 10kb bins.

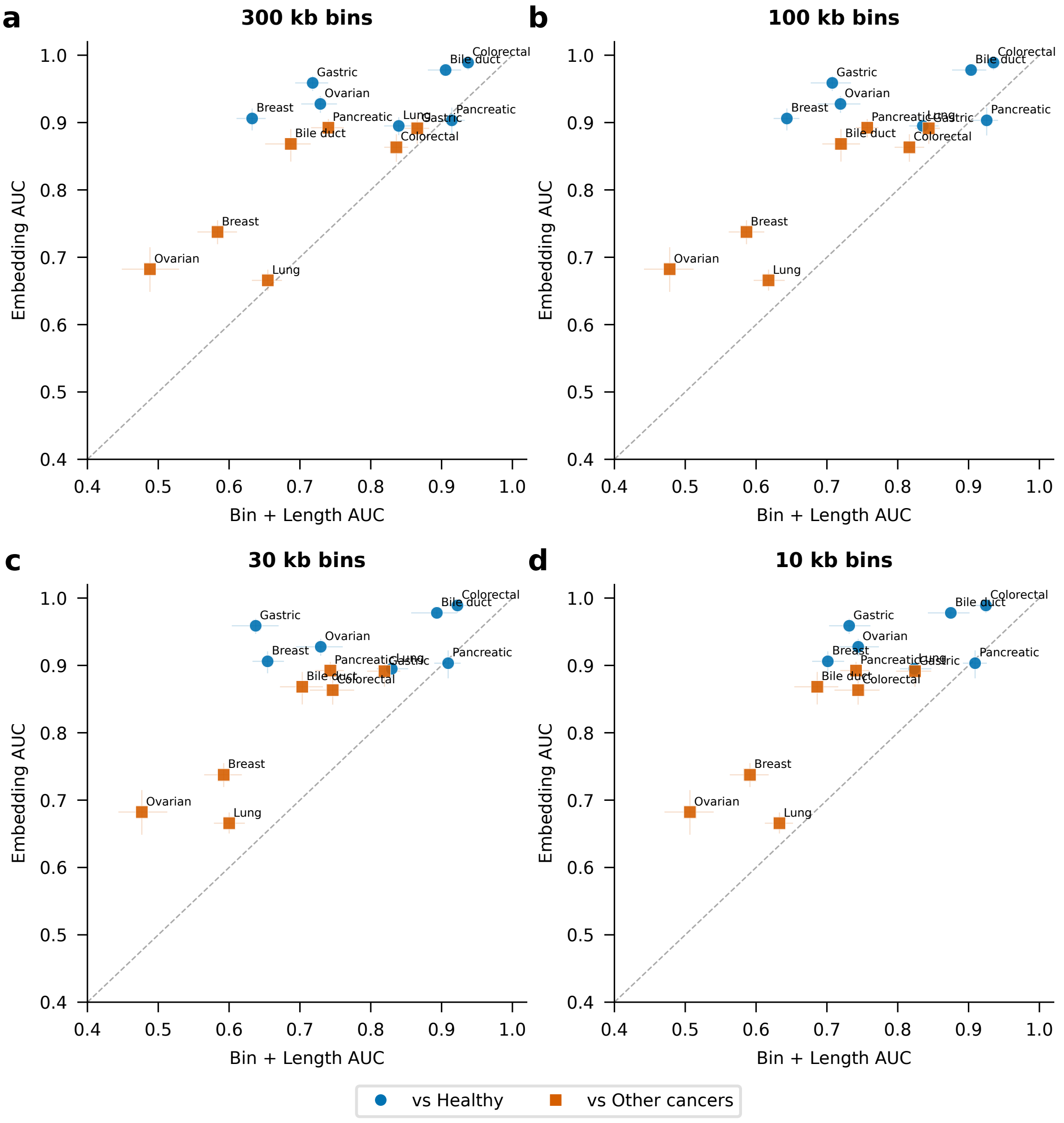


**Supplementary Figure 10.** LEAF-1+MIL embeddings versus Bin + Length features across bin resolutions in cfDNA. Per-cancer AUC for LEAF-1+MIL embeddings (y-axis) versus Bin + Length coordinate features (x-axis) in the Cristiano cohort[2], evaluated under matched cross-validation (see **Methods**). Each point is one cancer type; blue circles show cancer-versus-healthy detection and orange squares show cancer-versus-other-cancers discrimination. Error bars indicate confidence intervals along both axes, and the dashed diagonal indicates equal performance between the two representations. **(a)** 300kb bins. **(b)** 100kb bins. **(c)** 30kb bins. **(d)** 10kb bins.

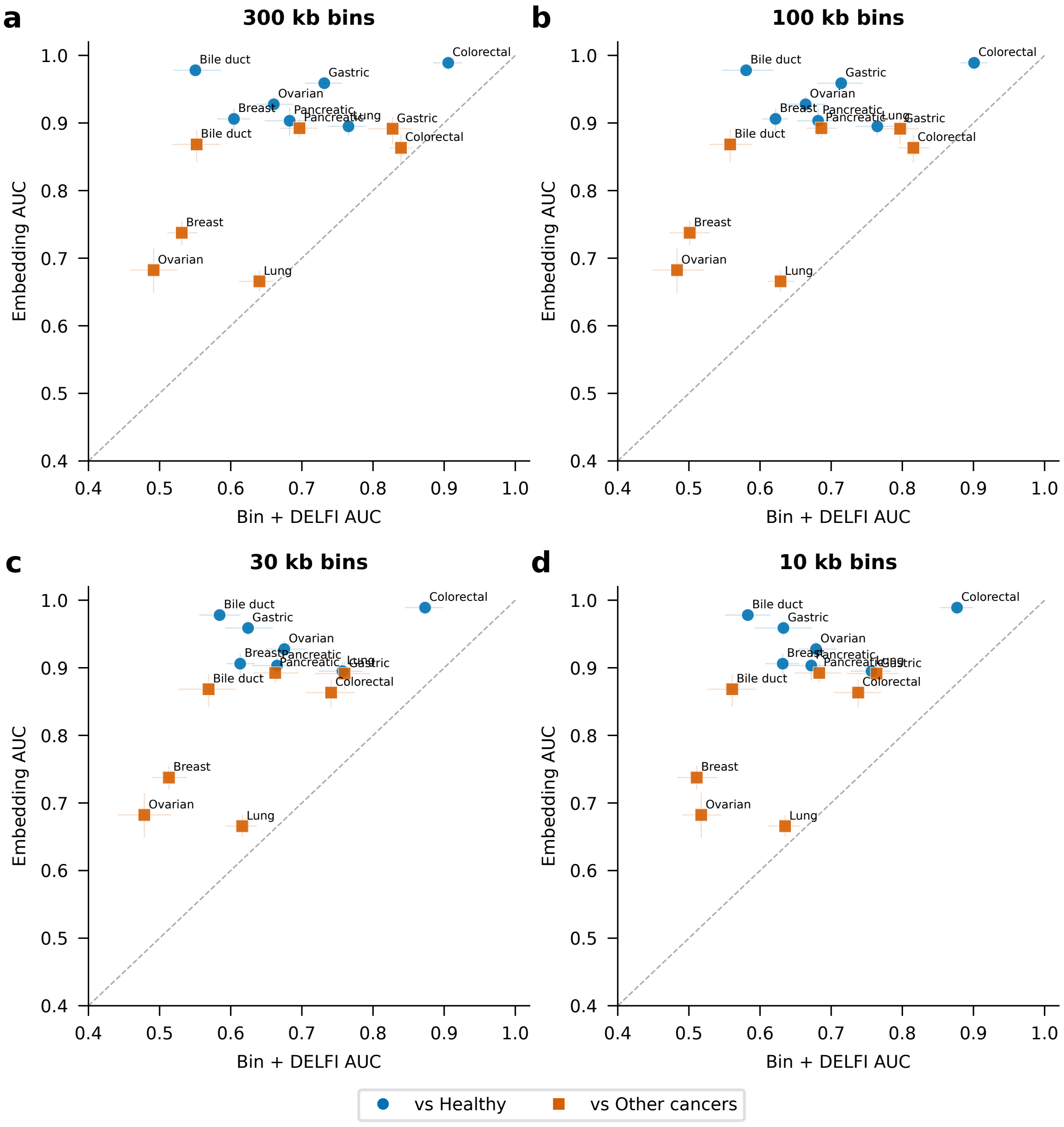


**Supplementary Figure 11.** LEAF-1+MIL embeddings versus Bin + DELFI features across bin resolutions in cfDNA. Per-cancer AUC for LEAF-1+MIL embeddings (y-axis) versus Bin + DELFI coordinate features (x-axis) in the Cristiano cohort[2], evaluated under matched cross-validation (see **Methods**). Each point is one cancer type; blue circles show cancer-versus-healthy detection and orange squares show cancer-versus-other-cancers discrimination. Error bars indicate confidence intervals along both axes, and the dashed diagonal indicates equal performance between the two representations. **(a)** 300kb bins. **(b)** 100kb bins. **(c)** 30kb bins. **(d)** 10kb bins.

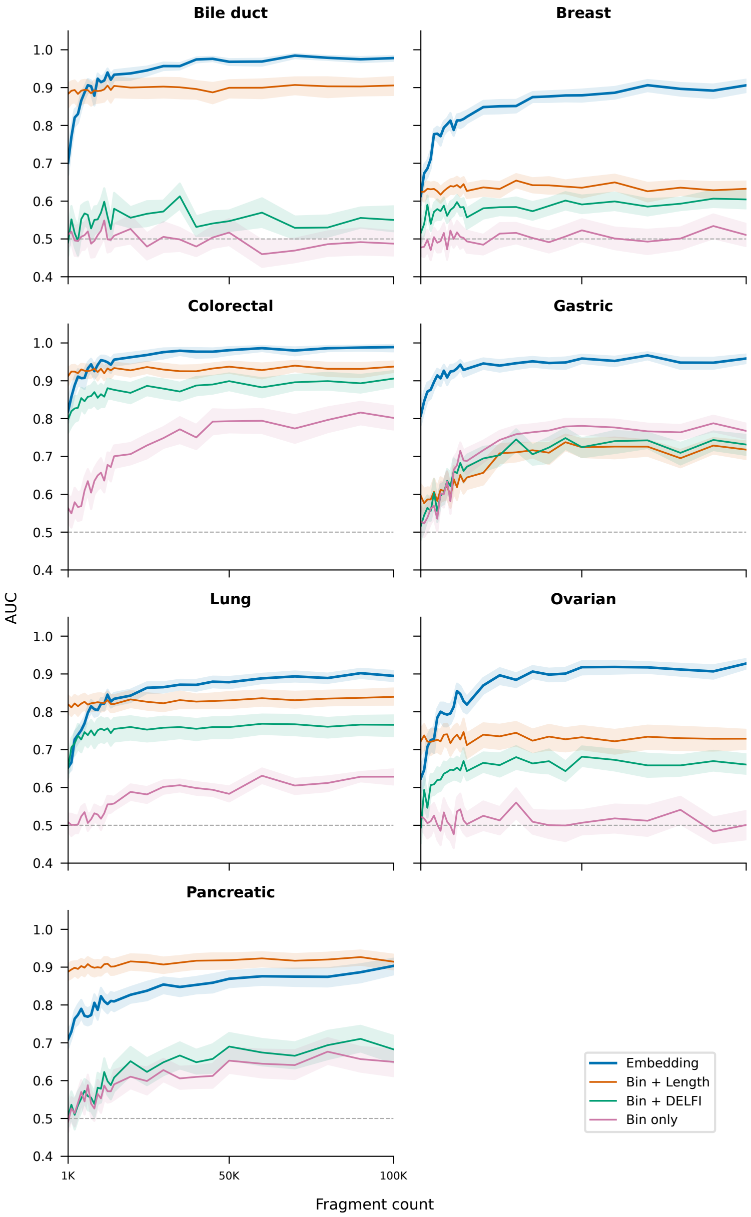


**Supplementary Figure 12.** Per-cancer cfDNA subsampling analysis: cancer versus healthy. Cancer-versus-healthy detection AUC for each of seven cancer types in the Cristiano cohort[2] as a function of the number of fragments sampled per sample (1,000 to 100,000), comparing LEAF-1+MIL embeddings with coordinate-based baselines (see **Methods**). Each panel corresponds to one cancer type (bile duct, breast, colorectal, gastric, lung, ovarian, and pancreatic). Lines show LEAF-1 embeddings (blue), Bin + Length (orange), Bin + DELFI (green), and Bin only (pink); shaded bands indicate confidence intervals. The grey dashed line indicates chance.

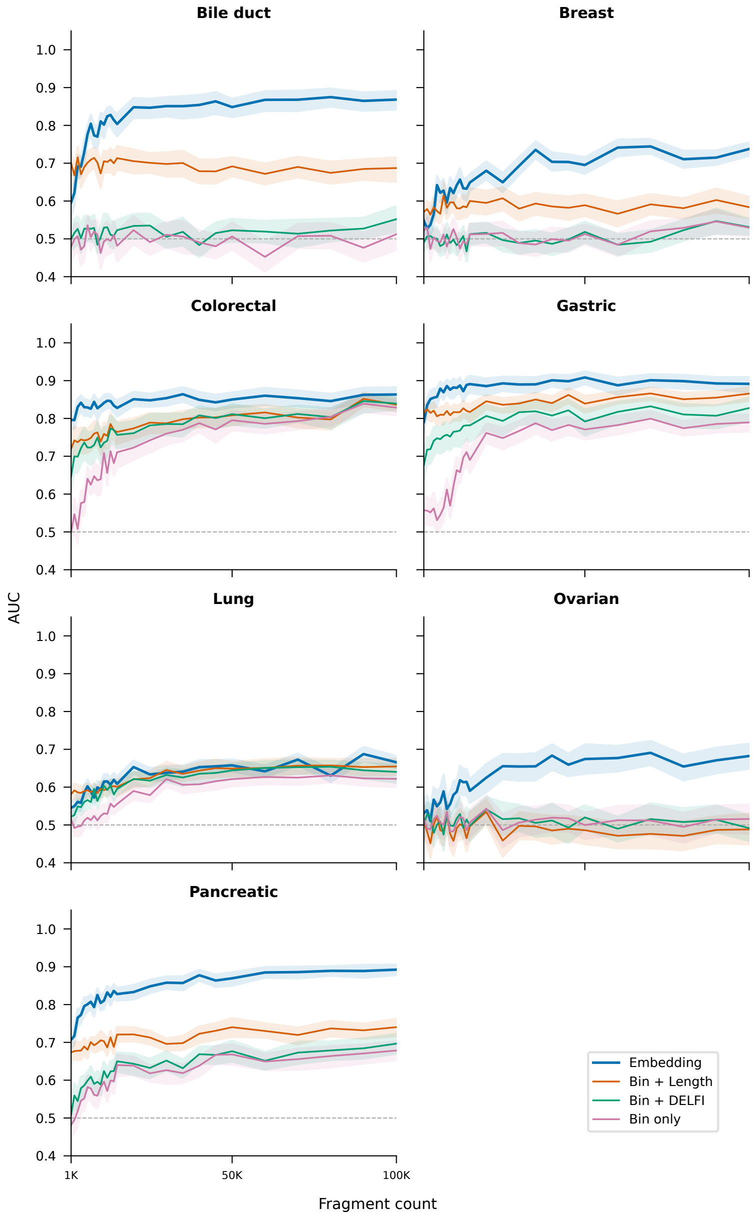


**Supplementary Figure 13.** Per-cancer cfDNA subsampling analysis: cancer versus other cancers (tissue of origin). Tissue-of-origin discrimination AUC (each cancer type versus the other cancer types) for each of seven cancer types in the Cristiano cohort[2] as a function of the number of fragments sampled per sample (1,000 to 100,000), comparing LEAF-1+MIL embeddings with coordinate-based baselines (see **Methods**). Each panel corresponds to one cancer type (bile duct, breast, colorectal, gastric, lung, ovarian, and pancreatic). Lines show LEAF-1 embeddings (blue), Bin + Length (orange), Bin + DELFI (green), and Bin only (pink); shaded bands indicate confidence intervals. Grey dashed line indicates chance.

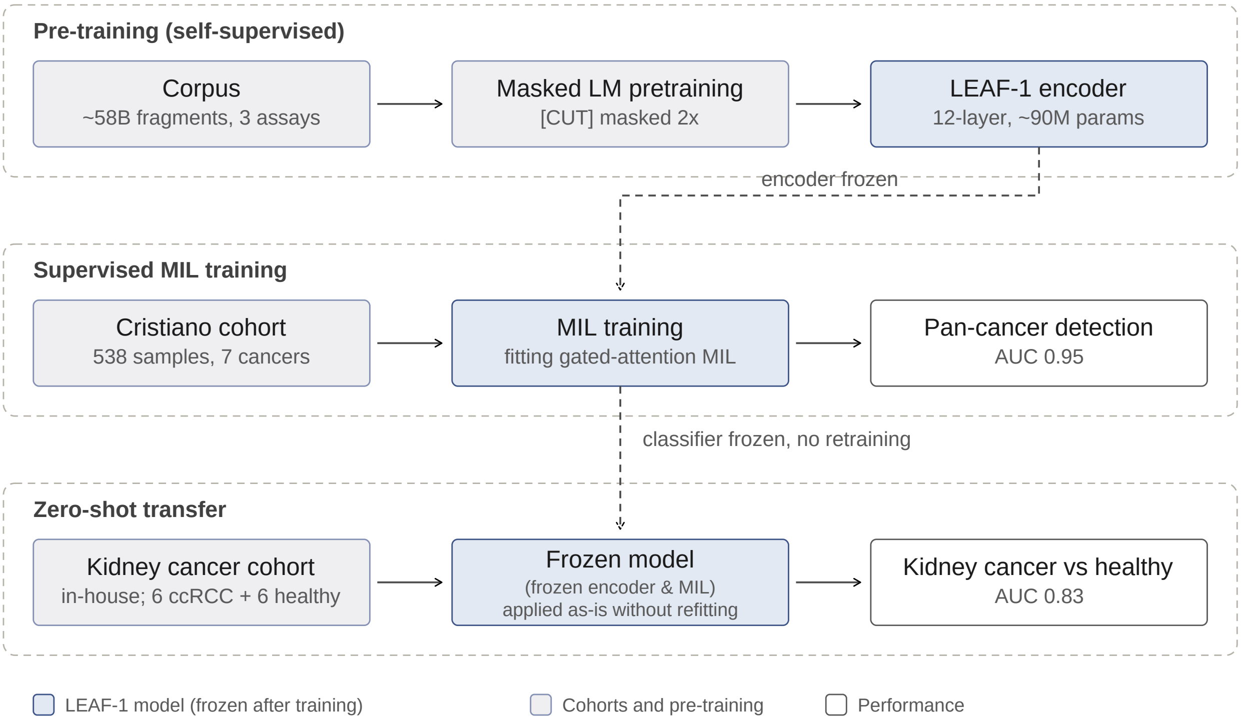


**Supplementary Figure 14.** Schematic of LEAF-1 pretraining, multiple-instance learning, and zero-shot transfer to an independent kidney cancer cohort. The LEAF-1 encoder (12-layer transformer, 90M parameters) was pretrained by masked language modeling on 58 billion fragments from cfDNA, bulk ATAC-seq, and single-cell ATAC-seq (2,620 samples; see **Methods**), with cleavage-boundary ([CUT]) tokens masked at twice the rate of sequence tokens. The frozen encoder (layer 4) was then used to embed 100,000 fragments per sample from the Cristiano cohort[2] (538 samples; seven cancer types and healthy controls), and a gated-attention multiple-instance learning (MIL) head was trained on these embeddings to produce a pan-cancer classifier (AUC 0.95). The frozen encoder and the frozen Cristiano-trained MIL classifier were then applied without retraining or threshold adjustment to an in-house cohort of six healthy donors and six patients with clear cell renal cell carcinoma profiled by shallow whole-genome sequencing, detecting kidney cancer at AUC 0.83. Dashed arrows indicate components reused without further training; box color denotes model components, classifier performance, and input cohorts, as shown in the legend.
